# A cortico-collicular circuit for accurate orientation to shelter during escape

**DOI:** 10.1101/2020.05.26.117598

**Authors:** Dario Campagner, Ruben Vale, Yu Lin Tan, Panagiota Iordanidou, Oriol Pavón Arocas, Federico Claudi, A. Vanessa Stempel, Sepiedeh Keshavarzi, Rasmus S. Petersen, Troy W. Margrie, Tiago Branco

## Abstract

When faced with predatorial threats, escape towards shelter is an adaptive action that offers long-term protection against the attacker. From crustaceans to mammals, animals rely on knowledge of safe locations in the environment to instinctively execute rapid shelter-directed escape actions ^1,2^. While previous work has identified neural mechanisms of escape initiation^3–5^, it is not known how the escape circuit incorporates spatial information to execute rapid flights along the most efficient route to shelter. Here we show that mouse retrosplenial cortex (RSP) and superior colliculus (SC) form a circuit that encodes shelter direction vector and is specifically required for accurately orienting to shelter during escape. Shelter direction is encoded in RSP and SC neurons in egocentric coordinates and SC shelter-direction tuning depends on RSP activity. Inactivation of the RSP-SC pathway disrupts orientation to shelter and causes escapes away from the optimal shelter-directed route, but does not lead to generic deficits in orientation or spatial navigation. We find that the RSP and SC are monosynaptically connected and form a feedforward lateral inhibition microcircuit that strongly drives the inhibitory collicular network due to higher RSP input convergence and synaptic integration efficiency in inhibitory SC neurons. This results in broad shelter direction tuning in inhibitory SC neurons and sharply tuned excitatory SC neurons. These findings are recapitulated by a biologically-constrained spiking network model where RSP input to the local SC recurrent ring architecture generates a circular shelter-direction map. We propose that this RSP-SC circuit might be specialized for generating collicular representations of memorized spatial goals that are readily accessible to the motor system during escape, or more broadly, during navigation when the goal must be reached as fast as possible.

---

Escaping to shelter has higher survival value than simply moving away from the source of threat^1^. Refuges are places where predators are usually impeded from entering and where predation risk is thus very low. To minimise exposure to the predator, navigating to the destination should in principle be as fast as possible, and animals rely on knowledge of the spatial environment to deploy accurate shelter-directed flights with very short reaction times ^1,2^. Mice in new environments can learn the location of a shelter within minutes, and when exposed to imminent threat orient their head and body in the shelter direction before running towards it along a straight vector^2^. Previous work has shown that shelter-directed escapes do not depend on sensory cues from the shelter and instead, mice use memory of the shelter location to reach to safety ^1,2^. Rodents therefore rely on continuously tracking of shelter direction during exploration to execute fast escape actions when exposed to threat^2,6^. Despite extensive knowledge about the neurobiology of spatial navigation^7–9^, it is not known how the escape circuit accesses information about safe locations to generate accurate shelter-directed actions during escape.

To investigate this problem, we allowed mice to explore a circular arena with a shelter and presented innately threatening sound stimuli ^3^. These threat stimuli reliably elicited shelter-directed escapes initiated with a head rotation movement that oriented the mouse in the shelter direction (Fig. 1A, Supplementary Video 1;^2^). To understand the neural basis of this orienting action we focused on two brain regions: the retrosplenial cortex (RSP), which has been shown to encode the direction and angular velocity of the head, as well as the spatial locations of goals and rewards^10^; and the superior colliculus (SC), which can generate orienting motor commands^11,12^ as well as escape initiation^3,13^, and receives projections from the RSP^14,15^. By simultaneously recording single unit activity in these two regions while the animal explored the arena (Extended Data Fig. 1A; Supplementary Video 2; 836 RSP units, 683 SC units, 4 mice) we found that the firing rate of a subset of neurons in both the RSP and SC (centro- and latero-medial regions) is tuned to shelter direction (Fig. 1B,C; Supplementary Video 3; Extended Data Fig. 1B,C). This information is encoded in an egocentric frame of reference and represents the angle between the current heading and the shelter (head-shelter angle; Supplementary Video 3). The preferred tuning of these shelter-direction neurons covers the entire egocentric angular space (Fig. 1C, Extended Data Fig. 1C, D) and their firing rate is independent from allocentric head direction (Extended Data Fig. 1E, Extended Data Fig. 2). To confirm that the firing fields of shelter-direction neurons are anchored to the shelter, we rotated its position by 90 degrees while keeping all other landmarks in place, which resulted in escapes to the new location after a brief exploration period^2^ (Fig. 1B; Supplementary Video 4). The firing fields rotated with the shelter location and therefore maintained their angular tuning profile with respect to the shelter position (RSP median rotation=113°, IQR=[83°, 134°]; SC median rotation=99°, IQR=[83°, 127°]; P>0.9 vs 90° and P=0 vs 0°, *non-uniformity v-test*; Fig. 1D), indicating that neurons in RSP and SC specifically and dynamically encode the shelter direction.

**Figure 1.**
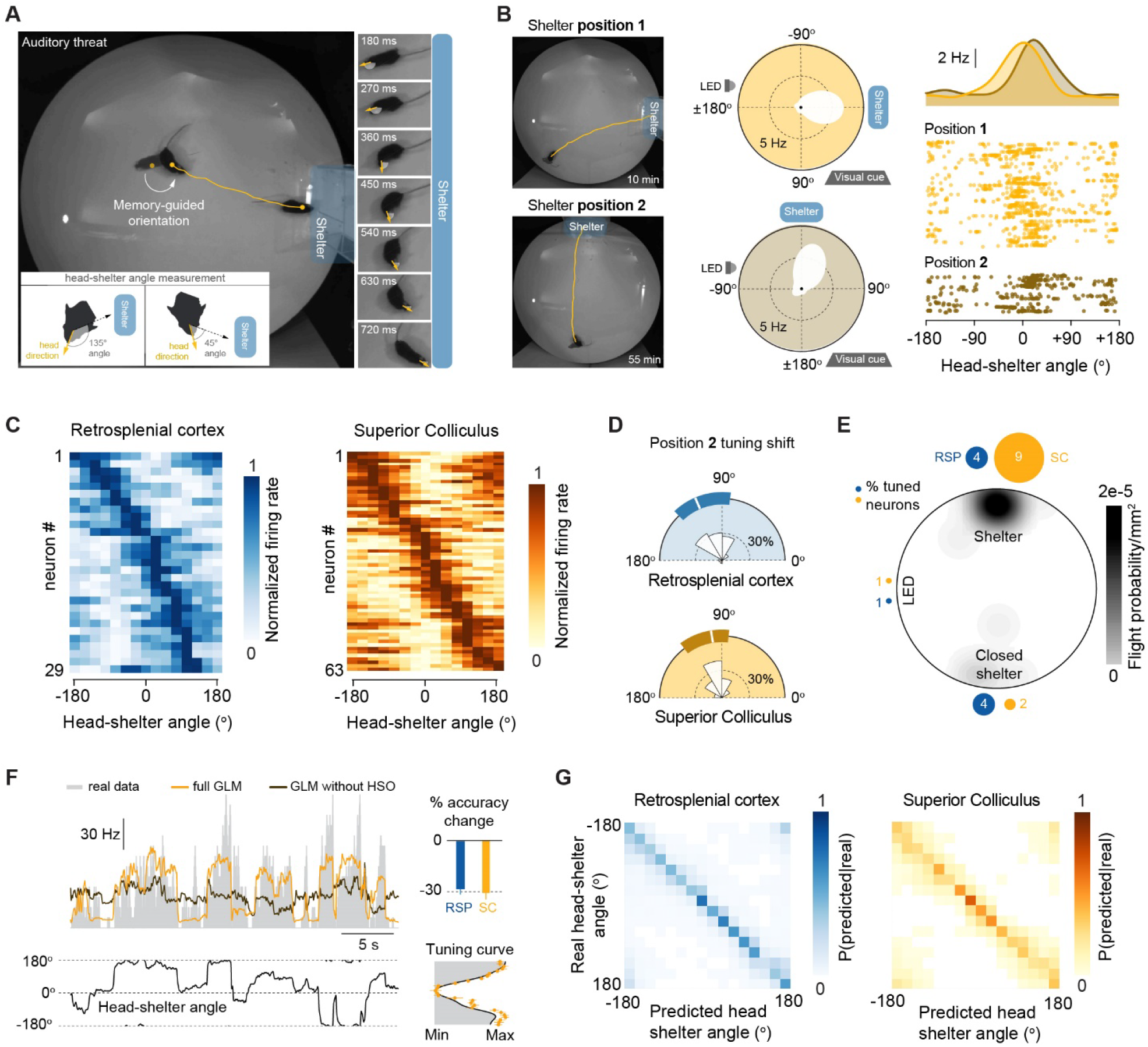
RSP and SC neurons encode egocentric shelter direction. **(A)** Left, superimposed video frames showing orientation to shelter after a threat stimulus and subsequent shelter-directed escape (yellow line shows flight trajectory); inset illustrates measurement of egocentric shelter direction (defined as the head-shelter angle). Right, video frame sequence detailing the decrease in head-shelter angle at escape initiation (yellow arrows: head direction, white shade: head-shelter angle). **(B)** Left, video frames with escape trajectories (yellow lines) to initial shelter position (top, position 1) and after rotation (bottom, position 2). Centre, tuning curves for a single SC unit, showing a rotation of the firing field that follows the rotation of the shelter position. Other explicit landmarks remain in place (LED and visual cue). Right, sample raster plot and tuning curve for the same neuron. **(C)** Tuning curves for shelter-direction RSP and SC units for the shelter in position 1. Curves sorted by tuning peak. **(D)** Distribution of change in the Rayleigh vector direction after rotating the shelter by 90° (position 1 to position 2). **(E)** Schematic of flight termination probability across the arena and percentage of units tuned to each of the shelters and the LED landmarks (circle area is proportional to percentage).**(F)** Example GLM data for a SC neuron showing the evolution of recorded firing rate (top-left) and head-shelter angle (bottom-left), together with the full model fit (cross validated prediction accuracy: 50±3%) and the same model without the head-shelter angle variable. Bottom right shows the tuning curve recovered from the full model fit (yellow symbols) and the recorded curve (gray). **(G)** Cross-validated confusion matrices for LDA population decoding of head-shelter angle.

The tuning of shelter-direction neurons could in principle reflect either a behaviourally relevant or perceptually salient location in the environment. To distinguish between these two possibilities, we added to the arena a second, identical shelter but with the entrance closed, as well as a bright LED. Mice preferentially explored the open and closed shelters’ locations (Extended Data Fig. 1F), and threat presentation resulted in 79% escapes to the open shelter, 14% to the closed shelter and none to the LED (N=29 trials; Fig. 1E), indicating the relative behavioural relevance of each location for escaping from imminent threat. We then computed the egocentric direction tuning to these two locations, as well as to the location of one the landmarks in the arena, a bright LED. While the fraction of RSP neurons tuned to the open and closed shelter was similar (open shelter: 3.5%, closed shelter: 3.5%; P=0.721, *two-proportions z-test*), SC neurons were preferentially tuned to the open shelter (open shelter: 9.2%, closed shelter: 1.6%; P=0.000638, *two-proportions z-test*), with both regions being significantly less sensitive to LED location (RSP: 0.8%, P=0.0277; SC: 1.1%, P=0.00254; *two-proportions z-test* vs open shelter; Fig. 1D). This suggests that, in this behavioural context, while the RSP contains a broader representation of familiar places in the arena, neurons in the SC preferentially encode the direction of the most behaviourally salient location for escape.

Neurons in the SC and RSP have been shown to encode a range of different behavioural variables^10,16^, so we next quantified the contribution of shelter tuning to the firing rate of shelter-direction neurons in both areas. We built a generalised linear model that included several variables, including head-direction and angular head movements (see Methods), and measured the difference in firing rate prediction accuracy between the full model and one without shelter direction (Fig. 1F). Removing the shelter-direction predictor caused a ∼30% drop in prediction accuracy, confirming that this variable is a key determinant of firing rate in single RSP and SC neurons (relative accuracy drop: RSP=-24±3.67%, P=1.2511e-06; SC=-31± 3.6%, P=8.8599e-12, *one-tailed t-test*). Furthermore, at the population level, a linear discriminant analysis-based decoder (LDA) could use the neuronal firing rates to predict the shelter direction significantly above chance (prediction accuracy: SC=0.73, RSP=0.83; chance=0.19; Fig 1G). Together these results show that there are neurons in the RSP and SC that represent the ongoing shelter direction in an egocentric reference frame, which is the information needed to generate orienting movements towards the shelter during escape.

As the RSP is proposed to act as a hub for integrating spatial information^10^, we hypothesised that the representation of shelter-direction in the SC might depend on RSP input. To test this, we assessed the effect of inactivating RSP neurons on SC shelter direction tuning. We used a chemogenetic approach where we targeted the inhibitory designer receptor hM4Di selectively to SC-projecting RSP neurons by injecting AAVretro-Cre^17^ in the SC and Cre-dependent hM4Di in the RSP (Fig. 2A and Extended Data Fig. 3). We then recorded single unit activity in the RSP and SC during activation of hM4Di with systemic delivery (i.p.) of CNO. This manipulation significantly decreased the *in vivo* firing rate of RSP neurons, confirming the chemogenetic inhibition (P=0.0029 *one-tailed Kolmogorov-Smirnov test*; N=340 units, 2 mice; Fig. 2B, C). In the SC, while the overall firing rate of the population was not affected (Extended Data Fig. 4), RSP loss-of-function caused a 33% decrease in the fraction of SC shelter-direction neurons (P=0.01, *Chi square test*; Fig. 2D). Accordingly, the ability to decode shelter direction at the SC population level was also compromised (Fig. 2E; median change in accuracy: −18.76%; IQR: [−15.7%, −19.7%], P=0.0020 *sign rank test*). These data show that the representation of egocentric shelter direction in the SC depends on RSP input. They further suggest that the RSP-SC pathway might be a critical neural circuit element for navigating to shelter during escape.

**Figure 2.**
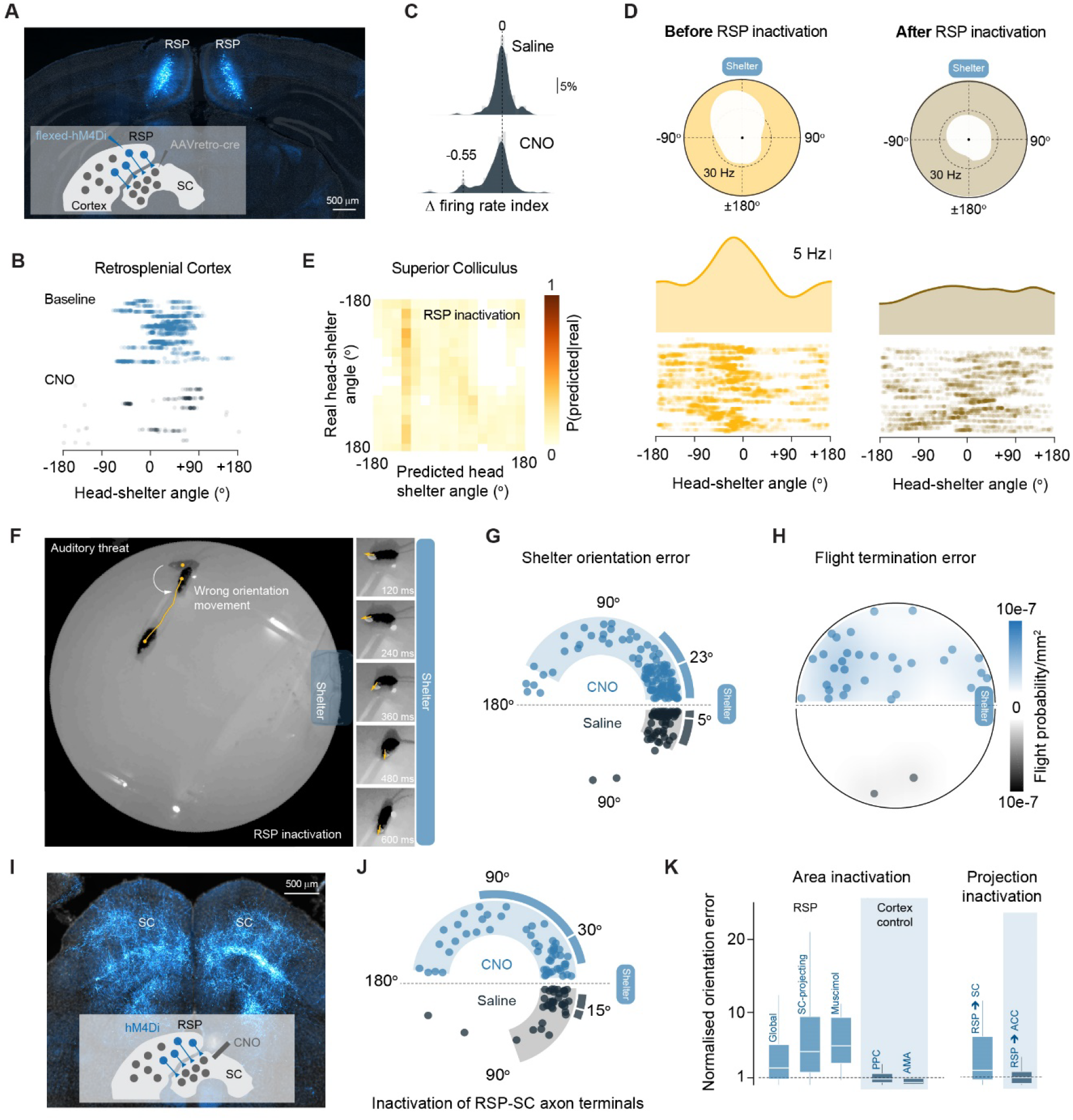
Shelter-direction tuning in SC and orientation to shelter during escape depend on RSP input. **(A)** Schematic of retrograde AAV strategy for inactivating SC-projecting RSP neurons, and coronal image of targeted RSP neurons. **(B)** Sample raster plot of an RSP shelter-direction cell before and after i.p. CNO. **(C)** Population histograms showing a decrease in firing rate in RSP neurons after i.p. CNO (see methods for index calculation) **(D)** Example tuning curves and sample raster plots showing loss of tuning in an SC shelter-direction neuron after i.p. CNO. **(E)** Cross-validated confusion matrix for LDA population decoding of head-shelter angle in SC after RSP inactivation. **(F)** Example video frames as in Fig. 1A showing escape initiation in the wrong direction due to incorrect orientation. **(G)** Summary plot of orientation errors at escape initiation after i.p. CNO. Light shaded area is 5-95 percentile range; dark bar is IQR and white line shows median. **(H)** Summary plot for location of flights terminated outside the shelter(CNO: 29.5%, saline: 3.9%). **(I)** Example coronal image of RSP axon terminals in the SC and schematic for local inactivation of these terminals. **(J)** Orientation errors as in B for local CNO infusion over SC (54 CNO trials and 37 saline trials from 6 mice; P=0.0032 permutation test). **(K)** Summary of activity manipulation effects on orientation errors. (Global RSP: 57 CNO trials and 32 saline trials from 5 mice, P=0.004; Muscimol: 21 muscimol trials from 6 mice and 25 saline trials from 3 mice, P=0.0061; PPC: 31 CNO trials and 22 saline trials from 3 mice, P=0.6514; AMA: 31 CNO trials from 4 mice and 28 saline trials from 3 mice P=0.1003; RSP to ACC: 21 CNO trials and 21 saline trials from 5 mice, P=0.8282;permutation test). Data are normalized by dividing each inactivation trial error by the median of orientation error of the respective control dataset.

To investigate the relevance of RSP neurons and their SC projections for shelter-directed escape behaviour, we used the same retrograde AAV strategy to inactivate SC-projecting RSP neurons (Extended Data Fig. 3) while monitoring behavioural responses to imminent threat. Loss-of-function of these RSP neurons caused mice to make larger errors in the orientation phase of escape (Fig. 2F, G; Supplementary Video 5; median escape error: CNO = 23°, 105 trials, 11 mice; saline=5°, 51 trials, 6 mice; P = 0.0153 *permutation test*) without affecting escape vigour (Extended Data Fig. 6G). This phenotype was observed for escapes elicited in both light and dark conditions, indicating the errors were not caused by impaired visual processing (Extended Data Fig. 5). The orientation deficits resulted in flight trajectories that were not directed towards the shelter location and consequently, the probability of terminating the flight outside the shelter was 7.4 times higher than the control (P=0.0001 *two-proportions z-test*; Fig. 2H). Although these escapes after RSP-SC inactivation were often resumed and the shelter eventually found, the time to shelter was 50% longer that observed for the control (CNO: median=2.1s, IQR=[1.8s, 2.8s]; saline: median=1.4s, IQR=[1.1s, 1.7s]; P=0.0439 *permutation test*; Extended Data Fig. 6H). Furthermore, flights were aborted earlier when orientation errors were largest (Extended Data Fig. 6I), suggesting that mice might retain awareness of shelter direction but are unable to select the appropriate direction of travel, similar to topographic disorientation in humans with RSP lesions^18^. We observed the same orientation impairment after global inactivation of RSP with chemogenetics (Supplementary Video 6; Extended Data Fig. 7A) and muscimol (Supplementary Video 7; Extended Data Fig. 7D). However, chemogenetic loss-of-function of the posterior parietal cortex (PPC) and anterior motor areas (AMA), two other SC-projecting cortical areas involved in goal directed navigation, did not perturb shelter-directed escapes (Fig. 2K, Extended Data Fig. 7B-C, Extended Data Fig. 8). This confirms that orienting to shelter is particularly dependent on RSP function. Since systemic CNO injection may inactivate SC-projecting RSP neurons that have collaterals to other regions, we next tested whether the orientation deficits were specifically due to loss of activity in RSP-SC synapses. We again targeted hM4Di to SC-projecting RSP neurons with the retrograde viral strategy, but locally infused CNO over the SC to specifically inactivate the RSP-SC projection^19^ (Fig. 2I; Supplementary Video 8; Extended Data Fig. 9A), which recapitulated the escape orientation errors of global RSP loss-of-function (Fig. 2J). In contrast, placing the cannula over a prominent secondary projection of SC-projecting RSP neurons, the anterior cingulate cortex (Extended Data Fig. 9B), did not affect escape behaviour (Fig. 2J; Supplementary Video 9), further supporting a unique role of RSP-SC inputs in orienting to shelter during imminent threat.

Since previous studies have shown that the SC commands orienting to sensory stimuli ^11,16 20^ and that the RSP contains representations of space ^8,10, 21^, the behavioural phenotype we observe after inactivation of the RSP-SC pathway could in principle trivially result from perturbing these known functions. To evaluate this possibility, we first tested whether RSP-SC inactivation impaired the classic role of the SC in sensory-guided orienting. We started by considering that RSP input to the SC might be necessary for SC control of head movement, and analysed whether RSP inactivation affected the relationship between the firing rate of SC neurons and head movements. As previously reported^22^, we found SC neurons tuned to egocentric head displacements (Fig. 3A), from which upcoming angular head movements could be decoded at the population level (Extended Data Fig. 10A). Chemogenetic inactivation of SC-projecting RSP neurons with i.p. CNO injection did not affect motor tuning of SC neurons (Fig. 3A) and the LDA prediction accuracy was not different between CNO and saline conditions (Fig. 3C; median change: 2.1%; IQR: [0.2%,3.2%], P=0.11 *sign rank test*). This suggests that SC motor function was not affected by RSP-SC inactivation, therefore predicting that simple orientation movements to acute sensory stimuli should be preserved after RSP loss-of-function ^11,16,20^. To test this prediction, we next developed a sound orienting assay to probe sensory-guided SC motor function. One of two speakers placed opposite each other emitted a brief tone while the animal explored the arena. Mice innately oriented to the sound emitting speaker quickly and as accurately as they orient to the shelter (Fig. 3B; Extended Data Fig. 10B-E; Supplementary Video 10). However, in contrast to the shelter orientation errors during escape, inactivation of SC-projecting RSP neurons (Extended Data Fig. 3) did not increase the error in orienting to the sound, confirming that RSP activity is not required for sensory stimulus-guided orientation (P=0.9290 *permutation test*; N=23 CNO trials and 26 saline trials, 4 mice; Fig. 3D; Supplementary Video 11). These experiments demonstrate that the escape errors after RSP-SC inactivation do not result from disrupting the general ability of the SC to command orientation movements. Instead they indicate that the RSP-SC pathway is specific for memory driven goal orientation, suggesting that the SC contains parallel streams for processing direct sensory orienting and orienting to high-order memory-based targets.

**Figure 3.**
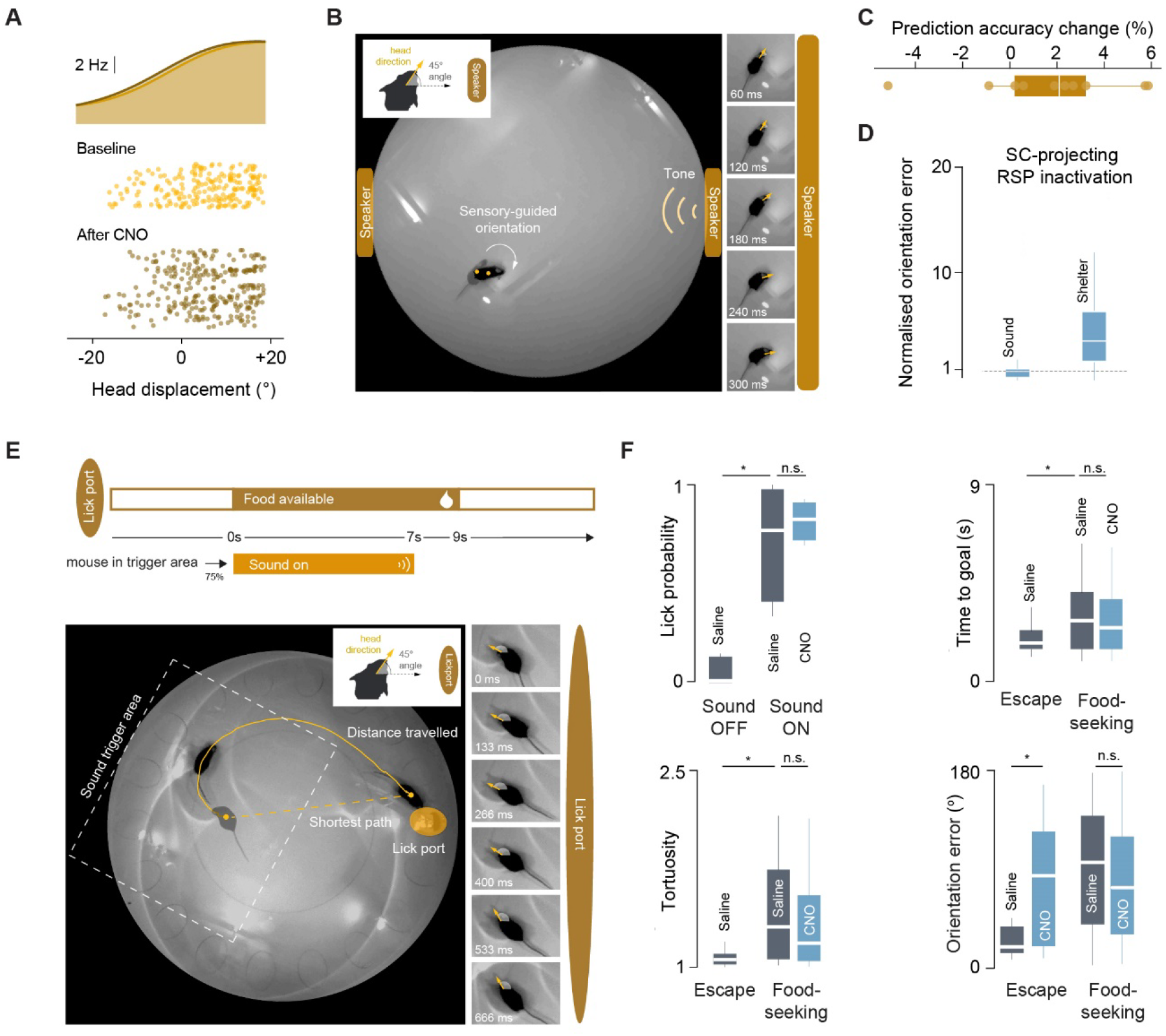
Orientation and navigation errors after RSP-SC inactivation are specific to escape behaviour. **(A)** Tuning curve and sample raster plot for an SC neuron tuned to egocentric head displacement. **(B)** Example video frames showing instinctive orientation to a sound source, an SC-dependent behaviour. **(C)** Change in LDA prediction accuracy of head displacement from SC firing rates (% change in CNO - % change in saline). **(D)** Summary data showing no effect of inactivation of SC-projecting RSP neurons in the sound-orienting assay but increased shelter orientation errors for the same mice (P=0.0143 permutation test; 32 CNO trials and 42 saline trials). Data are normalized by dividing each CNO trial error by the median of orientation error of the respective control dataset. **(E)** Task schematic (top) and example video frames showing curved trajectory and slow orientation towards lick port during a learned, non-urgent goal navigation task. **(F)** Summary data showing that mice learned the task rules (top left) and that the behaviour is not affected by inactivation of SC-projecting RSP neurons - lick probability, time to lick port, trajectory tortuosity (defined as the ratio between distance travelled by mouse and shortest possible path) and errors in orientation to lick port are the same for saline control and CNO trials. This contrasts with the increased shelter orientation errors during escape measured in the same mice. Time to goal and path tortuosity are different between navigation during escape and food-seeking. ^*^: p-value<0.05.

We next addressed whether escape errors after RSP inactivation arise because of a generic disruption of the ability of mice to navigate in space. First, we analysed navigation during exploratory behaviour and found no differences between control and RSP-SC inactivated animals (Extended Data Fig. 6 E-F). Second, we probed the effect of RSP-SC inactivation on goal-directed navigation in an alternative task. To test this, we placed mice in the same arena, but instead of a shelter we provided a lick port to which the mice were trained to go to in response to a long sound cue (7s). Although both navigation to the lick port and navigation to the shelter during escape are directed towards a learned, rewarding spatial goal, a key difference in the navigation strategies in these two tasks was the efficiency with which the mice reach the goal: mice arrived at the lick port ∼1s slower than at the shelter during escape, and the path taken to the lick port was curved, unlike the direct path taken during escape, as the mice did not orient immediately to the target but instead slowly oriented throughout the trajectory (Fig. 3E, F). In contrast with escape, inactivation of RSP-SC did not increase orientation errors to lick port (Fig. 3F; Supplementary Video 12). These data show that the escape errors following RSP-SC inactivation do not simply reflect a generic deficit in spatial navigation. Instead, they suggest that the RSP-SC pathway might have a specialized role for either escape or navigation when the goal must be reached as quickly as possible, and similar to the SC, suggest that there are parallel pathways in the RSP network for different types of goal directed navigation ^10,23^.

To understand the cellular and circuit mechanisms supporting the transfer of shelter direction information between the RSP and the SC we next measured the dynamics of the SC network in response to RSP input using single unit recordings in head-fixed animals and optical stimulation via a fibre placed over the SC (Fig. 4A and Extended Data Fig. 14). We expressed ChrimsonR, a red-shifted opsin, in SC-projecting RSP cells to activate RSP input, and channelrhodopsin-2 in either vGluT2^+^ or vGAT^+^ cells in the SC to determine the response profile of excitatory and inhibitory SC neurons, respectively, using an optotagging approach^24^ (Fig. 4A). Activation of RSP input to SC with 20 Hz pulses caused a short-latency increase in the firing rate of both excitatory and inhibitory SC neurons (vGAT^+^ latency: 9 ms, vGluT2^+^ latency: 4 ms; Fig. 4B). Over the course of 1000 ms of stimulation, however, the firing profiles of these two cell populations diverged markedly: while the firing rate of vGAT^+^ neurons increased progressively, the activity of vGluT2^+^ neurons started to decrease after ∼100 ms and became inhibited, falling below the baseline firing rate. This low-dimensional activity profile captures most of the variance across SC neurons responses to cortical activation (Extended Data Fig. 11).While this stimulation paradigm is artificial, it indicates that the RSP engages both excitatory and inhibitory cells in SC. In addition, the divergent firing profiles between the two SC populations suggested to us that there might be a feedforward lateral inhibition circuit, whereby inhibitory cells activated by RSP inhibit excitatory cells that also receive RSP input. We therefore investigated in more detail the functional organization of the RSP-SC microcircuit.

**Figure 4.**
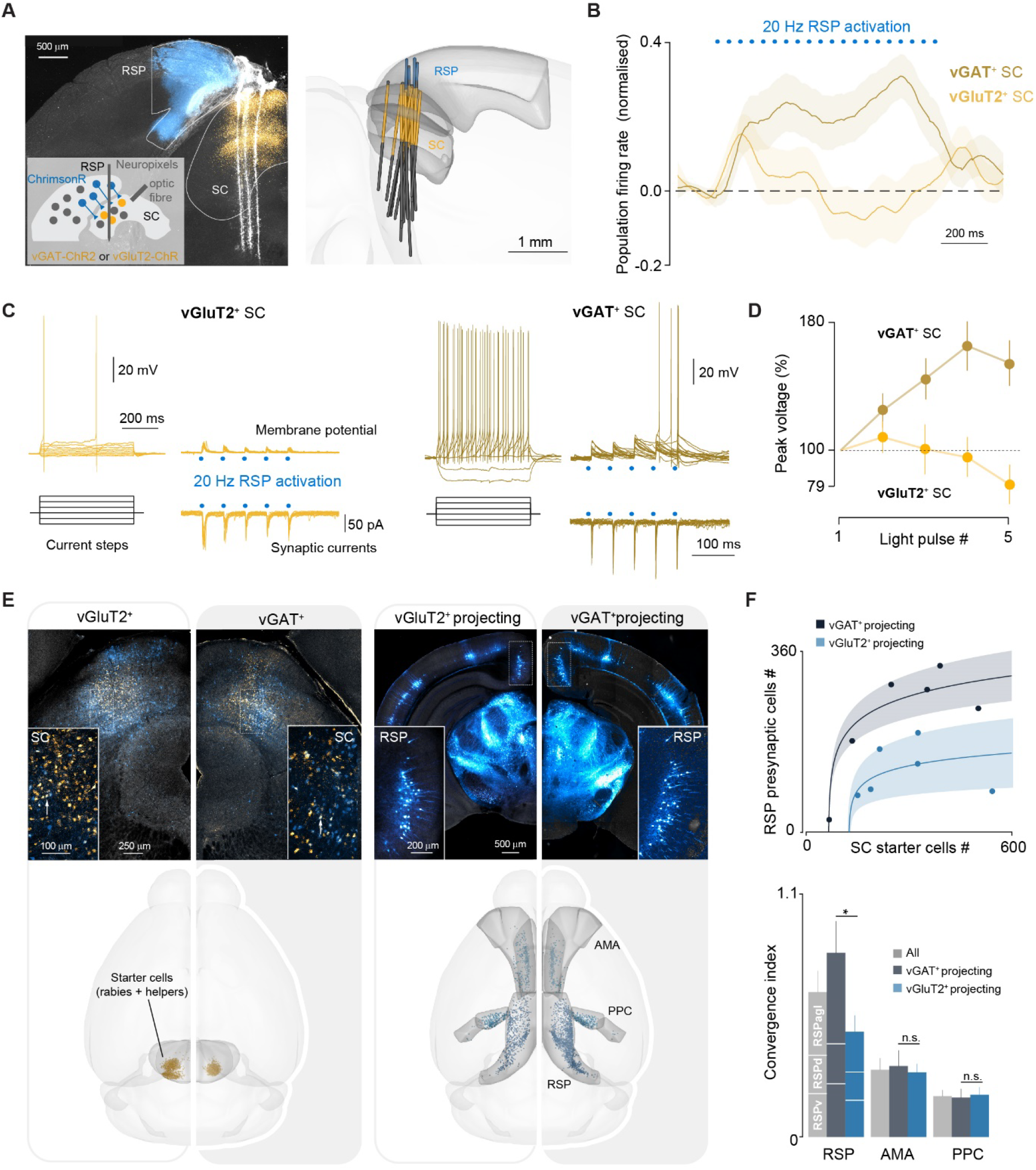
RSP synaptic input has a diverging effect on SC vGAT^+^ and vGluT2^+^ neurons. **(A)** Schematic of dual opsin strategy for recording dynamics of the SC network following RSP input in head-fixed mice (inset) and coronal image of the targeted RSP and SC neurons with trajectories of acutely inserted Neuropixels probes (left). Right, 3D rendering of probe trajectories in all mice. **(B)** Normalised firing rate of vGluT2^+^ (32 neurons) and vGAT^+^ (68 neurons) SC populations during 20 Hz activation of RSP axons in SC *in vivo*. Both SC populations are initially activated but diverge in firing profile over the course of stimulation. **(C)** Example synaptic currents and potentials evoked by 20 Hz optogenetic stimulation of RSP inputs onto excitatory and inhibitory SC neurons *in vitro* (blue circles indicate stimulation times). Left insets show voltage response to step current injections. **(D)** Summary plot of summation during 20 Hz optogenetic activation of RSP inputs (vGAT^+^: slope=17.6%, P=5.2e-13; VGIuT2^+^: slope=-2.9%, P=0.16, linear regression). **(E)** Left, coronal images (top) and 3D reconstruction (bottom) of vGluT2^+^ or vGAT^+^ starter cells in SC. Cells expressing helper AAVs are shown in yellow, rabies-infected cells in blue, and starter cells in white (inset) White arrows point to example starter cells. Right, coronal image (top) and 3D reconstruction (bottom) showing labelled input areas to excitatory and inhibitory SC neurons, including the RSP (inset). **(F)** Top, number of RSP labelled cells as a function of the number of SC starter cells, showing higher convergence of RSP input onto inhibitory SC cells for any given number of starter cells (each dot represents data from one mouse). Fitted functions: logarithmic; fitting p-values 0.0016925 vGlut2^+^ projecting and 1.7e-5 vGAT^+^ projecting. Shaded area represent 95% confidence interval. Note that the confidence intervals for vGlut2^+^ projecting and vGAT^+^ projecting never overlap. Bottom, cell type-specific convergence index for RSP, AMA and PPC, showing that the higher convergence onto SC inhibitory neurons is specific to RSP inputs and that AMA and PPC have lower convergence onto SC neurons overall. Starter cells were observed in all the three RSP subdivisions: RSP ventral part (RSPv), RSP dorsal part (RSPd) and RSP lateral agranular part (RSPa).

First, we characterised the anatomy of the RSP-SC projection^14,15^ with cell type-specific monosynaptic retrograde rabies tracing^25^ from vGluT2^+^ and vGAT^+^ starter cells (Fig. 4E; N=6 each mice). Rabies infection of either SC population resulted in prominent labelling of layer 5 RSP neurons, showing that both excitatory and inhibitory SC cells indeed receive RSP input (Fig. 4E). There was, however, 1.8 times more convergence of RSP input onto inhibitory than excitatory cells (Fig. 4F; convergence index vGAT^+^: 0.83±0.14, vGluT2^+^: 0.47±0.08, P=0.035 *t-test*). The magnitude of RSP convergence into the SC was also higher than two other cortical areas involved in navigation (PPC and AMA), which projected in a sparser and symmetrical manner to excitatory and inhibitory SC neurons (AMA: convergence index vGAT^+^: 0.29 ± 0.05, vGluT2^+^: 0.32 ± 0.05; PPC: convergence index vGAT^+^ 0.19 ± 0.04, vGluT2^+^ 0.18 ± 0.04, Fig. 4F). Next, we measured the physiological properties of RSP-SC connections using whole-cell recordings *in vitro* and channelrhodopsin-2 expressed in the RSP. Optogenetic activation of RSP axons over the centromedial and centrolateral SC confirmed a high connectivity rate for both excitatory and inhibitory SC neurons (VGluT2^+^: 37.1%, N=70 cells, 4 mice; vGAT^+^: 43.8%, N=73 cells, 4 mice). RSP input elicited excitatory monosynaptic potentials in both cell populations, and we found that temporal summation was more efficient in vGAT^+^ neurons because of differences in intrinsic excitability and short-term synaptic plasticity (Fig. 4C, D; Extended Data Fig. 12; 5^th^ pulse peak voltage: 154±14% for vGAT^+^, 79±12% for vGluT2^+^, P=2.53e-4 *t-test*). Together, these data show that both excitatory and inhibitory SC neurons receive monosynaptic input from the RSP, and suggest that RSP activity should generate a stronger drive in the inhibitory network because of higher anatomical convergence and more efficient synaptic integration in vGAT^+^ neurons.

Next, we looked for the existence of the feedforward lateral inhibition motif suggested by the RSP stimulation experiments, using a dual opsin, dual recombinase anterograde strategy and slice electrophysiology. Briefly, to express an opsin specifically in the SC vGAT^+^ cells that receive RSP input, we injected an anterograde virus carrying a Cre-dependent flippase ^26^ into the RSP of vGAT::Cre mice. This anterograde virus jumps one synapse into the SC, resulting in SC vGAT^+^ cells expressing flippase. Injection of a flippase-dependent channelrhodopsin-2 into the SC then produced opsin expression in this cell population. In addition, we also expressed a Cre-off fluorescent marker to label putative excitatory SC cells (i.e.: vGAT negative), as well as ChrimsonR in the RSP to activate RSP axons. We then obtained whole-cell recordings from SC excitatory cells, activated ChrimsonR to identify vGAT-negative neurons that directly receive RSP input and stimulated channelrhodopsin-2 to elicit inhibition. This experiment revealed that 26.3% of excitatory neurons in the SC receive both monosynaptic input from the RSP and are directly inhibited by the local vGAT^+^ neurons that also receive RSP input (Fig. 5A-C). The RSP-SC circuit therefore contains a feedforward lateral inhibition motif.

**Figure 5.**
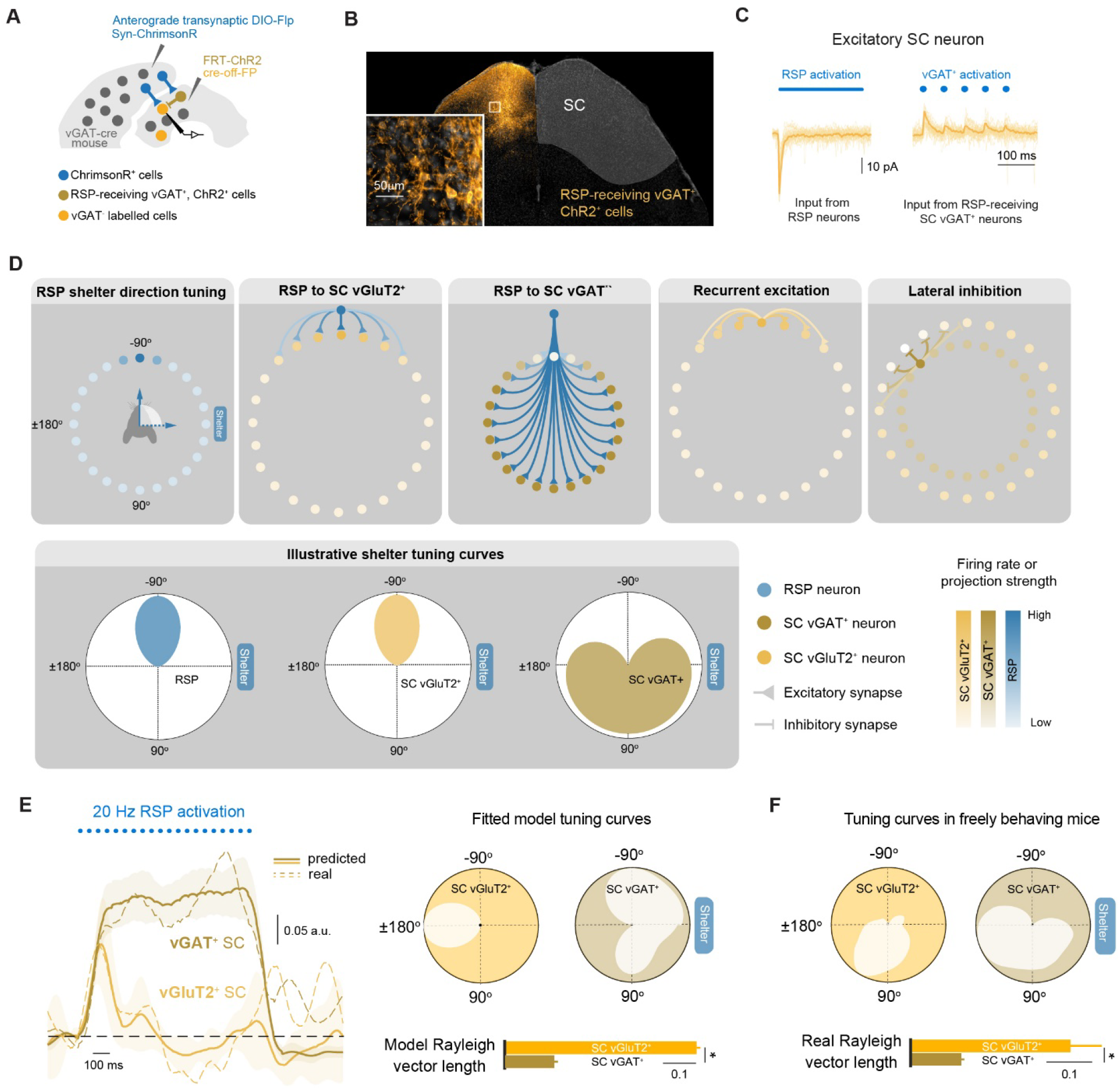
A feedforward lateral inhibition model for mapping shelter direction from RSP to SC. **(A)** Schematic of the anterograde dual opsin strategy used to test for feedforward lateral inhibition *in vitro*. **(B)** Coronal image of RSP-receiving vGAT^+^ cells expressing ChR2. **(C)** Activation of RSP-receiving vGAT^+^ SC cells generates inhibitory synaptic currents in vGATSC cells (right) that also receive monosynaptic RSP input (left). **(D)** Top: Schematic illustrating the ring architecture and connectivity profiles for the different components of the feedforward lateral inhibition model (RSP, vGluT2^+^ SC and vGAT^+^ SC networks). Neurons are shown organized in a circular fashion with respect to their shelter tuning preference. Colour intensity of circles indicates neuronal firing rate, colour intensity of lines indicates projection strength. From left to right: RSP population firing when the mouse faces −90 degrees from the shelter, RSP projections to vGluT2^+^ SC and vGAT^+^ SC (only projections from the RSP neuron turned to −90 degrees are shown), local recurrent excitation and lateral inhibition within SC. RSP neurons excite a small number of recurrently connected SC vGlut2^+^ neurons, which inherit the same tuning, while exciting all other SC vGAT^+^ neurons that then project to orthogonal vGlut2^+^ SC neurons. Activity in a given RSP neuron results in feedforward lateral inhibition of all SC vGlut2^+^ neurons with a different shelter direction preference. Additional circuit elements are shown in Extended Data Fig 13. Bottom: tuning curves schematics for a RSP neuron tuned to −90 degrees and for the vGluT2^+^ and vGAT^+^ SC neurons most strongly excited by the RSP neuron shown. **(E)** Left: mean and 95% confidence interval of predicted firing rate of vGluT2+ and vGAT+ SC populations following 20 Hz activation of RSP neurons in the best 30 models fitted to the data, compared to the dynamics recorded in vivo (dashed lines, also in 4B). Right: example predicted tuning curves for vGluT2+ and vGAT+ SC neurons (top). The model predicts that vGluT2+ SC neurons are more sharply tuned to shelter direction than vGAT+ SC neurons (bottom). **(F)** Experimentally measured tuning curves of opto-tagged vGluT2^+^ and vGAT^+^ SC neurons and Rayleigh vectors lengths for the population of recorded neurons

To further explore how the RSP-SC circuit maps shelter direction between RSP and SC, we measured the electrophysiological properties of additional local SC connections (Extended Data Figure 13A) and built a spiking network model constrained by all our anatomical and electrophysiological data. Briefly, the model consisted of conductance-based leaky integrate-and-fire RSP neurons organised as a ring-like network, with neighbouring neurons tuned to similar angles and providing inputs to vGluT2^+^ and vGAT^+^ SC neurons with diagonal and inverted diagonal connectivity kernels, respectively (Fig. 5D; see experimental procedures for details). vGAT^+^ SC neurons inhibited vGluT2^+^ SC neurons to replicate the measured feedforward lateral inhibition architecture, and the model also included the remaining local SC connections, all with diagonal connectivity matrices (Fig. 5D and Extended Data Fig. 13 B). This model successfully fitted the SC firing dynamics elicited by RSP stimulation (average sum of squared errors of 30 best fitting models, 1.6770 ± 0.0066; Fig. 5E), while a standard ring attractor model without feedforward lateral inhibition^9,27–29^ achieved a poorer fit (4.1891 ± 0.0743, t-test p-value 8.97*10^−40^; Extended Data Figure 13C). In the feedforward lateral inhibition model, when the animal is at a particular angular distance from the shelter, RSP neurons tuned to this distance create an activity bump in a small fraction of excitatory SC neurons, which inherit the respective cortical tuning (Fig. D). In parallel, the active RSP neurons also excite SC inhibitory neurons that project to an orthogonal population of excitatory SC cells, and consequently silence the excitatory neurons surrounding the activity bump because of the high efficiency of the RSP synaptic connection onto inhibitory SC neurons (Fig. 5E). The net result is a clean mapping of shelter direction between the RSP and SC networks. A prediction from this model is that the tuning of SC excitatory cells should be narrow, whereas the tuning of inhibitory cells should be broad because they pool from RSP neurons with diverse tuning. To test this prediction, we recorded SC shelter-direction cells *in vivo* while optotagging excitatory or inhibitory cells (Extended Data Fig. 1A and 14). We found that the tuning curves of inhibitory SC neurons were indeed broader than for excitatory cells, with Rayleigh vectors 2.4 times higher (vGAT^+^: 0.09 ± 0.002, vGluT2^+^: 0.22 ± 0.04, t-test 0.0054; Fig. 5 F).

Our results show that the RSP and SC continuously encode the escape goal direction vector in an egocentric reference frame. RSP neurons project to both excitatory and inhibitory SC neurons but generate a stronger drive of the inhibitory network because of higher convergence and more efficient synaptic integration. By dissecting the RSP-SC pathway with projection and cell-type specificity, we find a feedforward lateral inhibition circuit architecture, whereby inhibitory neurons that receive RSP input directly inhibit excitatory neurons that also integrate RSP synaptic input. Since the representation of shelter direction in the SC depends on RSP activity, a plausible model is that the RSP generates an egocentric representation of shelter direction that the SC inherits. While it is possible that the RSP broadcasts spatial and self-motion signals that are used locally by the SC to assemble shelter direction *de novo*, we show that the RSP-SC synaptic connection is necessary for accurate orientation to shelter during escape. We therefore favour a model where shelter direction is directly mapped from the RSP to the SC. Through simulating a biologically-constrained spiking network we propose that this mapping is done by exploiting the feedforward lateral inhibition motif to generate shelter tuning in a subset of excitatory SC neurons while inhibiting the network of excitatory cells with orthogonal tuning. The finding that shelter tuning curves in excitatory SC neurons are sharper than in inhibitory neurons is in support of this model. Similar models have been previously applied to explain tuning specificity in sensory pathways, including visual^30^, somatosensory^31^ and auditory^32^; as well as to the representation of saccadic vectors between the frontal eye field and the SC^33^. Our model is also closely related to models of heading direction in a variety of circuits and animals species^27,34^, perhaps suggesting a conserved circuit architecture for mapping circular variables.

The firing fields of shelter-direction cells are similar to goal-vector cells found in the hippocampus of bats^35^ and the lateral entorhinal cortex of rats navigating to learned reward locations, and to egocentric center-bearing cells in the postrhinal cortex; though distinct from egocentric boundary vector cells^21,36^. The finding of an egocentric representation of shelter direction vector in the RSP agrees with previous work showing that RSP neurons encode behaviourally important locations, such as landmarks, reward locations and a variety of spatial features of the environment^21,37^. In particular, the RSP integrates information from multiple streams to form representations of head-direction and angular velocity, place, and conjunctives of these variables^21^, and it is thus well-positioned to compute spatial representations in egocentric coordinates^10,36,38^. An important finding of our study is that while the escape orientation errors after RSP-SC inactivation could in principle be explained by a perturbation of these known RSP functions and a general loss of navigation ability, we show that loss-of-function in the RSP-SC pathway does not perturb navigation during exploration or to a food reward goal. This suggests that there are parallel circuits for different types of navigation, and that the RSP-SC pathway might be dedicated for escape behaviour or navigation when the goal needs to be reached as fast as possible. Our recordings with two shelters in the arena further suggest that the RSP may form egocentric maps of behaviourally important locations in general. In contrast, the SC specifically encoded shelter direction. Together with the known role of the SC in establishing saliency and priority maps^39,40^, these findings raise the possibility that the SC selects the most relevant goal-relevant target from a pool of RSP-encoded locations.

Previous work has demonstrated a role for cortico-collicular pathways mediating learned actions in highly trained animals in a variety of tasks and species^41,42^. Here we demonstrate that a cortico-collicular circuit is essential for an instinctive, survival behaviour. A key finding is that SC motor function and sensory-guided orientation were not affected by RSP loss-of-function, and thus escape orientation errors cannot be explained by generic orientation deficits. Instead, our data suggest a selective role for the RSP-SC circuit in the memory-guided orienting movements required to orient to shelter ^2^. Future work will be needed to understand how the shelter orienting action is generated upon the detection of threat, which requires integrating additional information from areas such as the medial SC^4^. A possible advantage of this cortico-collicular organization is to use cortical circuits to perform complex computations and distil the result to variables that can be easily converted into actions – here, the shelter direction continuously mapped already in egocentric coordinates. This could potentially decrease the time to execute the accurate action, which in the case of escaping from imminent threat is of great survival value. The model that emerges from our results may therefore represent a generic brain strategy for using cortical output to generate fast and accurate goal-directed actions.

## Experimental procedures

### Mice

All experiments were performed under the UK Animals (Scientific Procedures) Act of 1986 (PPL 70/7652 and PFE9BCE39) following local ethical approval. Male adult C57BL/6J wild-type (Charles Rivers, 6 – 24 weeks old) were used for all behavioural experiments. Mice were single housed after surgery and at least 3 days before starting behavioural protocols. Mice had free access to food and water on a 12:12 h light:dark cycle and were tested during the light phase (unless stated otherwise). VGluT2::eYFP (resulting from in-house cross of VGluT2-ires-Cre with R26-stop-eYFP, JLS #016963 and #006148 respectively; JLS: Jackson Laboratory stocks) or VGAT::eYFP (resulting from in-house cross of VGAT-Cre with R26-stop-eYFP, JLS #016962 and #006148 respectively) mice (5 – 8 weeks old) were used for ChR2-assisted circuit mapping (CRACM). VGluT2-ires-Cre (JLS #016963) or VGAT-Cre (JLS #016962) mice (9 – 20 weeks old) were used for retrograde rabies tracing and dual opsin-assisted circuit mapping and opto-tagging experiments.

### Surgical procedures

Mice were anaesthetized with an intraperitoneal (i.p.) injection of ketamine (95 mg kg^−1^) and xylazine (15.2 mg kg^−1^) or with isoflurane (2.5% - 5%) and secured on a stereotaxic frame (Kopf Instruments).. Isoflurane (0.5–2.5% in oxygen, 1 l min^−1^) was used to maintain anaesthesia. Carprofen (5 mg kg^−1^) was administered subcutaneously for analgesia. Craniotomies were made using a 0.5 mm burr and coordinates were measured from bregma or lambda (see Supplementary Tables 1 and 2 for implant affixation and virus injection coordinates, respectively). Viral vectors were delivered using pulled glass pipettes (10 μl Wiretrol II pulled with a Sutter-97) and an injection system coupled to a hydraulic micromanipulator (Narishige), at approximately 50 nl min^−1^. Implants were affixed using light-cured dental cement (3M) and the surgical wound was sutured (Vicryl Rapide) or glued (Vetbond).

### Viruses

The following viruses were used in this study and are referred to by contractions in the text. Chemogenetic projection-specific inhibition experiments: rAAV2-retro-CMV-bGlo-iCre-GFP (made in house; 1.07×10^12^ GC ml^-1^; ^17^) and AAV2-hSyn-DIO-hM4D(Gi)-mCherry (Addgene #44362; 4.6×10^12^ GC ml^-1^; ^19^); chemogenetic non-projection specific inhibition experiments AAV8-hSyn-hM4D(Gi)-mCherry (Addgene #50475; 4.8×10^12^ GC ml^-1^). Retrograde rabies tracing^25^: EnvA pseudotyped SADB19 rabies virus (SAD-B19^ΔG^-mCherry; made in house; 2×10^8^ GC ml^-1^) used in combination with AAV2/1 encoding the EnvA receptor TVA and rabies virus glycoprotein (AAV2/1-Flex-N2cGnucGFP; made in house; 7×10^12^ GC ml^-1^ and AAV2/1-EF1a-Flex-GT-GFP; made in house; 2×10^12^ GC ml^-1^); Retrograde rabies tracing for automated cell counting (see Anatomical tracing): EnvA pseudotyped N2c (N2c^ΔG^ - mCherry; made in house; 10^9^ GC ml^-1^) used in combination with a single AAV2/2 helper virus encoding both the EnvA receptor TVA and rabies virus glycoprotein (AAV2/2-EF1a-(ATG-out)Flex-EBFP-T2a-TVA-E2A-N2cG, made in house; 1.13×10^13^ GC ml^-1^; ChR2-assisted circuit mapping: AAV2/1-CAG-hChR2(H134R)-mCherry-WPRE-SV40 (Penn Vector Core; 2×10^14^ GC ml^-1^) or AAV2/2-EF1a-DIO-hChR2(H134R)-mCherry (UNC Gene Therapy Vector Core; 5.1×10^12^ GC ml^-1^). Dual opsin-assisted circuit mapping and opto-tagging in head-fixed and freely moving mice: AAV2/-EF1a-DIO-hChR2(H134R)-EYFP -WPRE-pA (UNC Gene Therapy Vector Core; 4.2×10^12^ GC ml^-1^); AAV2/-Syn-ChrimsonR-tdT^43^ (Addgene plasmid 59171, 1.3×10^13^ GC ml^-1^). Dual opsin-assisted circuit mapping and opto-tagging in brain slices: AAV2/-Syn-ChrimsonR-tdT (Addgene plasmid 59171, 1.3×10^13^ GC ml^-1^); AAV2/1-CAG-Flex-FlpO (made in house; 2×10^12^ GC ml^-1^); AAV2/1-Ef1a–fDIO-hChR2(H134R)-EYFP (made in house; 9.2×10^14^ GC ml^-1^); AAV2/1-CamKIIa-Cre-OFF-GCaMP7f (made in house; 1.0×10^14^ GC ml^-1^).

### Behavioural procedures

#### Experimental setup

All behavioural experiments were performed in an elevated circular Perpex arena (92 cm diameter), located in an opaque, sound-deadening 140 × 140 × 160 cm enclosure^2^. The shelter was an over-ground red translucent Perspex box (20 × 10 × 10 cm) placed at the edge of the arena, with a 4.2 cm wide entrance facing the centre of the arena. The enclosure held six infrared light-emitting diodes (LED) illuminators (Abus) and experiments were recorded with an overhead near-IR GigE camera (Basler), at 30 or 40 frames per second. Unless stated otherwise, experiments were conducted at a background luminance of 2.7 lux, generated by a projector (BenQ) pointed at a translucent overhead screen (Xerox). Video recording and sensory stimulation (see below) were controlled with custom software written in LabVIEW (2015 64-bit, National Instruments) and Mantis software (mantis64.com). All signals and stimuli, including each camera frame, were triggered and synchronised using hardware-time signals controlled with a PCIe-6351 and USB-6343 board (National Instruments).

#### Sensory stimuli

##### Escape behaviour assay

The stimulus was a frequency-modulated auditory upsweep from 17 to 20 kHz over 2 s ^44^. In a small fraction of experiments additional stimuli were used to minimise habituation of escape responses. Waveform files were created in MATLAB (Mathworks), and the sound was generated in LabVIEW, amplified and delivered via an ultrasound speaker (Pettersson) positioned 50 cm above the arena centre. Sound pressure level at the arena floor was 73 - 82 dB.

##### Food seeking assay

same as before, but the stimulus that was a 7s tone at a frequency of 5 kHz..

##### Orientation to sound assay

The stimulus was a 300 ms tone at a frequency of 2.5 kHz. Sound pressure measured at the centre of the arena was 75 – 80 dB.

#### Behavioural protocols

##### Escape behaviour assay

the procedure was conducted as previously described ^2^. Before the experiment, bedding from the animal’s home cage was placed inside the shelter. After a 7 min habituation period, during which the mouse had to enter the shelter at least once, sound stimuli were delivered. During a single session, multiple stimuli were delivered with an inter-stimulus interval of at least 90 s. Typically, half of the trials were presented in the dark (0.01 – 0.03 lux); escape error was not different between dark and light conditions and therefore the results were pooled (P>0.05 for all datasets *permutation test*). Experiments had a typical duration of 90 minutes.

##### Orientation to sound assay

experiments were conducted in the dark using the same arena as the escape behaviour assay. The stimulus was delivered by one of two speakers at the height of arena floor, 10 cm away from the edge and positioned at diametrically opposite sides. After a 7 min habituation period, 7 stimuli were presented with an inter-stimulus interval of at least 90 s from a randomly selected speaker. Stimuli were typically presented when mice subtended an angle smaller than 120° towards the speaker because sound location in azimuthal plane has been shown to be unreliable above this range^45^. Experiments had a typical duration of 40 minutes.

##### Food-seeking assay

experiments were conducted using the same arena as the escape behaviour assay. Food-restricted mice were trained for 21 days in navigating to a lick-port dispensing condensed-milk, upon the presentation of a sound cue signalling reward availability. Reward delivery was triggered by licks. The sound cue was trigger only when the mouse was exploring a specific region of the arena floor (sound trigger area, Fig. 3E). The lick port was positioned opposite from the trigger area, in a shallow hole near the edge of the arena, which the mouse could access via a step. During training, the temporal duration of the sound cue and the trigger zone were progressively decreased. On the experiment day, reward was available during the presentation of the 7 s long sound cue, which was automatically triggered with a probability of 0.75, after the mouse entered the trigger zone. Reward remained accessible for 2 s after stimulus offset. Licking was detected as described in Campagner, Evans et al. 2019^46^.

#### Analysis

Behavioural video and tracking data were sorted into peri-stimulus trials and manually annotated. Detection of the stimulus was assed as previously described ^3^.

##### Escape behaviour assay

Onset of escape was determined by visual inspection of the video recordings and considered as either the onset of a head-rotation movement or an acceleration (whichever occurred first) after the mouse detected the stimulus^3^. Flight termination was defined as the first time the mouse stopped outside the shelter, re-oriented or arrived to shelter, after having initiated an escape. Escape error was computed as the shelter direction at the end of the orientation phase of escape (see Fig. 1A for example of an accurate orientation phase where escape error is close to 0°). Shelter direction was measured manually using a custom-designed Python-based graphic interface and varied between 0° (mouse heading perfectly aligned to the centre of the shelter entrance) and ±180° (positive angles are measured in the clockwise direction from the shelter, negative angles are measured in the anti-clockwise direction). Flight distance was computed as the ratio of the Euclidean distance between the mouse position at escape onset and at the flight termination, over the Euclidean distance between the mouse position at escape onset and the shelter entrance.

##### Food-seeking assay

*lick probability* was computed as the fraction of trials in which the mouse licked at least once while reward was available. *Time to goal* is the time it took the mouse to reach the lick port hole, starting from stimulus onset.

##### Orientation to sound assay

Orientation error was computed as the head– speaker angle at the end of the first orientation movement after the auditory stimulus presentation.

### Pharmacological inactivation

Mice were bilaterally implanted with guide and dummy cannulae (Plastics One) over RSP and given at least 96 h for recovery. On experiment day, the animals were infused with 1.0 – 1.2 μL Muscimol-BODIPY-TMR-X (0.5mg/ml, ThermoFisher) or vehicle per hemisphere and tested in the escape behaviour assay. Mice were anesthetised and internal cannulae projecting 0.5 mm below guide cannulae, were inserted and sealed with Kwik-Sil. Muscimol or vehicle were then infused at a rate of 150–200 nl min^−1^ using a microinjection unit (Hamilton, 10μl syringe; in Kopf Instruments Model 5000) connected to the internal canula through tubing (Plastics One) and a plastic disposable adaptors (Plastics One). Mice were given 35 minutes to recover before starting the escape behaviour assay.

To confirm infusion site, immediately upon termination of the behavioural assay, mice were anaesthetized with isoflurane (5%, 2 l min^−1^) and decapitated. The brain was sectioned coronally (100 μm) with a vibrotome (Campden Instruments) in ice-cold PBS (0.1 M), directly transferred to 4% paraformaldehyde (PFA) solution, and kept for 20 min at 4 °C. The slices were then rinsed in PBS, counter-stained with 4 ′,6-diamidino-2-phenylindole (DAPI; 3 μM in PBS), and mounted on slides in SlowFade Gold (Thermo Fisher) before imaging (Zeiss Axio Imager 2) on the same day.

### Chemogenetic inactivation

Mice were injected with a retro-AAV (rAAV2-retro-CMV-bGlo-iCre-GFP) into the SC and AAV2/2-hSyn-DIO-hM4D(Gi)-mCherry into the RSP for projection-specific inactivation, or with AAV2/8-hSyn-hM4D(Gi)-mCherry into the target locations for global inactivation. After at least 4 weeks mice were intraperitoneally injected with CNO (final concentration of 10 mg kg^-1^; Hellobio CNO freebase) or saline, during brief (<2 min) isoflurane anaesthesia (2 – 4% in oxygen, 1 L min^-1^). Mice were given 35 minutes to recover before starting behavioural experiments. Saline and CNO sessions for a given mouse were spaced in time at least 3 days. In a subset of the above mice, cannulae were implanted either in the SC or in the anterior cingulate cortex (ACC) to inactivate specifically the respective RSP projection^19^. Guide and dummy cannulae (Plastics One) were implanted into the target location at least 4 weeks after viral injection, and at least 4 days before behavioural experiments. Cannulae implantation and infusion were performed as described in the *Pharmacological inactivation experiment* section. CNO was diluted in buffered saline containing (in mM): 150 NaCl, 10 D-glucose, 10HEPES, 2.5 KCl, 1 MgCl2 and to a final concentration of 10 μM. 0.8 – 1.2 μL of CNO were infused.

### Anatomical tracing

Injections for monosynaptic rabies tracing^25^ from unilateral VGluT2^+^ or vGAT^+^ SC cells were performed with an angled approach to avoid infection of the ipsilateral RSP. In the first surgery, a mix of AAV1-Flex-N2cGnucGFP24 and AAV2/1-EF1a-Flex-GT-GFP25, or AAV2/2-EF1a-(ATG-out)Flex-EBFP-T2a-TVA-E2A-N2cG only, was injected in the left SC (10° ML angle; AP: −0.40 mm from lambda; ML: −1.00 mm; DV: −1.75 mm; injection volume = 20 −25 nL). SAD-B19DG-mCherry rabies virus, or N2c-mcherry,was injected vertically 5 days after the first procedure (same target location; injection volume = 25 – 30 nL). Mice were perfused 10 days after the second procedure. Brains were imaged by serial micro-optical sectioning 2-photon tomography^47^ (40 μm thick coronal sections; voxel size ML 2 μm, DV 2 μm, AP 5 μm). For cell counting, images were first registered to the Allen Mouse Brain Atlas (c 2015 Allen Institute for Brain Science. Allen Brain Atlas API. Available from: brain-map.org/api/index.html). mCherry-expressing and EBFP expressing cells in SC, AMA, RSP and PPC were automatically counted using the deep learning algorithm CellFinder^48^, manually curated and visualised using brainrender^49^. Starter cells were defined as cells co-expressing helper EBFP and rabies mCherry. Convergence index for a given presynaptic brain region was computed as the ratio between the number of mCherry-labelled cells and the total number of SC starter cells. One sample brain was sectioned using a cryostat (Leica 3050 S) and imaged with an epifluorescence microscope (Zeiss Axio Imager 2).

### *In vitro* whole-cell recordings

#### Preparation of acute midbrain slices

For CRACM experiments, male and female VGluT2::EYFP or VGAT::EYFP mice were injected with AAV2/1-CAG-hChR2(H134R)-mCherry-WPRE-SV40 in RSP or AAV2/2-EF1a-DIO-hChR2(H134R)-mCherry in SC. For dual-opsin assisted circuit mapping and and opto-tagging experiments, VGAT::cre mice were injected with AAV2/-Syn-ChrimsonR-tdT and AAV2/1-CAG-Flex-FlpO in RSP, and AAV2/1-Ef1a–fDIO-hChR2(H134R)-EYFP and AAV2/1-CamKIIa-Cre-OFF-GCaMP7f in SC.

After 2 weeks (or 4 weeks for dual opsin-assisted circuit mapping and opto-tagging experiments) mice were killed by decapitation following isoflurane anaesthesia. Brains were quickly removed and immediately immersed in ice-cold slicing solution containing (in mM): 87 NaCl, 2.5 KCl, 26 NaHCO_3_, 1.25 NaH_2_PO_4_, 10 glucose, 50 sucrose, 0.5 CaCl_2_, and 3 MgCl_2_, with an osmolarity of 281-282 mOsm, and constantly bubbled with carbogen (95% O_2_ and 5% CO_2_) for a final pH of 7.3. Acute coronal slices of 250 μm thickness were prepared at the level of the SC (−0.8 to 0.2 mm from lambda) using a Vibratome (Leica VT1200 S). Slices were collected and transferred to a recovery chamber containing slicing solution and stored under submerged conditions at near-physiological temperature (35°C) for 30 minutes, constantly bubbled with carbogen (95% O_2_ and 5% CO_2_). Slices were then transferred to a different recovery chamber and submerged in artificial cerebrospinal fluid (aCSF) solution containing (in mM): 125 NaCl, 2.5 KCl, 26 NaHCO_3_, 1 NaH_2_PO_4_, 10 glucose, 2 CaCl_2_, and 1 MgCl_2_, with an osmolarity of 293-298 mOsm and constantly bubbled with carbogen (95% O_2_ and 5% CO_2_) for a final pH of 7.3. Slices were further incubated at room temperature (19-23 °C) for at least 30 more minutes prior to electrophysiological recordings.

#### Recording electrodes

Pipettes were pulled from standard-walled filament-containing borosilicate glass capillaries (Harvard Apparatus, 1.5 mm OD, 0.85 mm ID) using a vertical micropipette puller (PC-10 or PC-100, Narishige) to a final resistance of 4-6 MΩ. Pipettes were backfilled with potassium methane sulfonate based solution containing (in mM): 130 KMeSO_3_, 10 KCl, 10 HEPES, 4 NaCl, 4 Mg-ATP, 0.5 Na_2_-GTP, 5 Na-Phosphocreatine, 1 EGTA, biocytin (1 mg mL^-1^), with an osmolarity of 294 mOsm, filtered (0.22 μm, Millex) and adjusted to pH 7.4 with KOH.

#### Data acquisition

Whole-cell patch-clamp recordings were performed with an EPC 800 amplifier (HEKA). Data were sampled at 25 kHz, low-pass Bessel filtered at 5 kHz, digitised with 16-bit resolution using a PCIe-6353 board (National Instruments) and recorded in LabVIEW using custom software. For recordings, slices were transferred to a submerged chamber and perfused with aCSF constantly bubbled with carbogen (95% O_2_ and 5% CO_2_). The solution was perfused at a flow rate of 2-3 ml/min with a peristaltic pump (PPS2, MultiChannel Systems or Minipuls 3, Gilson) and temperature was kept at 32-34°C. Cells were visualised with oblique illumination on an upright SliceScope Pro 1000 (Scientifica) using a 60x water-immersion objective (Olympus) or with DIC illumination on an upright SliceScope Pro 6000 (Scientifica) using a 40x water-immersion objective (Olympus).

For CRACM experiments, expression of ChR2-mCherrywas assessed based on fluorescence from mCherry expression using LED illumination (pE-100, CoolLED) at a wavelength of 565 nm. Target cells were identified based on fluorescence from EYFP expression using LED illumination (pE-100, CoolLED) at a wavelength of 490 nm. ChR2 was activated with wide-field 490-nm LED illumination (pe-100, CoolLED) using a train of five 1-ms long pulses at 20 Hz (maximum light intensity = 6 mW). The resting membrane potential was determined immediately after establishing the whole-cell configuration and experiments were continued only if cells had a resting membrane potential more hyperpolarized than −45 mV. For dual opsin-assisted circuit mapping and opto-tagging experiments, target putative vGluT2^+^ cells were identified based on the fluorescence from GCamp7f expression using 470nm wide-field LED illumination (pE-800, CoolLED) filtered by a 466/40nm bandpass filter (FF01-466/40-25, Semrock). ChR2 was activated with wide-field 435-nm LED illumination (pe-800, CoolLED) filtered by a 430/40nm bandpass filter (86349, Edmund Optics) using a train of five 1-ms long pulses (maximum light intensity = 13.5 mW), that was preceded by a 250ms long pulse of 635-nm wide-field LED illumination (pe-800, CoolLED)^43^ filtered by a 624/40nm bandpass filter (67035, Edmund Optics), maximum light intensity = 13 mW,. ChrimsonR was activated with wide-field 635-nm LED illumination (pe-800, CoolLED) filtered by a 624/40nm bandpass filter (67035, Edmund Optics) using a train of five 1-ms long pulses (maximum light intensity = 13 mW).

Input resistance (*R*in) and series resistance (*R*s) were monitored continuously throughout the experiment, and *R*s was compensated in current-clamp recordings. Only cells with a stable *R*s < 40 MΩ were analysed. Upon termination of the recording, the anatomical location of the recorded neuron within the slice was recorded using a 5x objective (Olympus) for future reference. After the experiment, slices were fixed with 4% PFA for 2 hours, kept in 4°C PBS overnight, and imaged using an epifluorescence microscope (Zeiss Axio Imager 2).

#### Analysis

Analysis was performed in python 3.7. Normalised peak voltages of EPSPs in a train were calculated by dividing the peak amplitude of each evoked EPSP to the first evoked EPSP. Membrane potential values stated in the text are not corrected for liquid junction potentials.

### Histological procedures

For general histology mice were anaesthetized with Euthatal (0.15–0.2 ml) and transcardially perfused with 10 ml of ice-cold PBS with heparin (0.02 mg ml−1) followed by 4% PFA in PBS solution. Brains were post-fixed overnight at 4 °C then transferred to PBS solution. Sections (50 μm) were cut with a cryostat (Leica CM3050S) and stained. Unless otherwise stated, sections were imaged with an epifluorescence microscope (Zeiss Axio Imager. Immunohistochemistry was performed to enhance the signal of hM4D-mCherry-expressing RSP neurons. Sections were initially blocked using a 5% normal donkey serum (NDS) in PBS solution. Subsequently, they were incubated overnight at 4°C with anti-RFP primary antibody (1:1000, rabbit; 600-401-379, Rockland), followed by a 2-hour incubation with Alexa-568 donkey anti-rabbit. Antibodies were diluted in 0.5% NDS and 0.05% Triton X-100. DAPI was used for counterstaining. Sections were mounted in SlowFade Gold (Thermo Fisher, S36936) onto slides before imaging.

### Single unit recordings in freely moving animals

#### Data acquisition

Mice were singly housed after probe implantation in a reversed 12 h light cycle and tested during the dark phase of the day light cycle. A single Neuropixels probe (phase3A, option1, 384 channels^50^) was chronically implanted in the RSP and SC. Before insertion, the probe shank was coated with DiI (1 mM in ethanol, Invitrogen) for track identification (Extended Data Fig. 1A). At the end of the experiment the mouse was perfused and the brain was imaged with an epifluorescence microscope (Zeiss AxioImager 2) or by serial micro-optical sectioning 2-photon tomography to confirm the location of probe implantation (see *Anatomical tracing* section). For single unit recording experiments paired with chemogenetic inactivation, probes were implanted in mice previously injected (4 weeks) with viruses as described in *Chemogenetic inactivation*.

Extracellular recordings in freely moving animals were performed in a similar behavioural apparatus as described in *Behavioural procedure*, with an over ground shelter (20 × 10 × 10 cm; 12 cm wide entrance) positioned by the edge and facing the centre of the platform. Two explicit visual cues were positioned in the environment: a yellow LED, always positioned on the ‘West wall’ of the behavioural cabinet, and an A2 white cardboard sheet, always positioned in the ‘South wall’ of the cabinet, both distal to the arena. A custom-made rotary joint (adapted from Doric AHRJ-OE_PT_AH_12_HDMI) was used to prevent the cables from twisting. Each mouse was tested multiple times with a minimum time interval between consecutive experiments of 3 days. During each session, the mouse explored the shelter in two locations (east pole of the arena: shelter position one; north pole of the arena: shelter position 2) for at least 30 minutes each. In a fraction of the experiments, after the mouse was tested in the protocol described above, the LED was moved to the east pole of the arena or a closed shelter was introduced at the south pole of the arena and then moved to the east pole. For Neuropixels recording experiments paired with chemogenetic inactivation, the intraperitoneal injection of CNO (or saline) was done during the behavioural experiment, after the mouse explored the shelter in the second position for at least 30 minutes. The mouse was not removed from the arena and was not anesthetised for the injection. Although the recording was never interrupted, the first 35 minutes following the injection were excluded from further analysis. Extracellular potentials were recorded with SpikeGLX (https://github.com/billkarsh/SpikeGLX, Janelia Research Campus) and were amplified (500 x), high-pass filtered (300 Hz) and sampled at 30 kHz.

#### Analysis

Data analysis was performed in MATLAB. Spike sorting was done using Janelia Rocket Cluster 3.0^51^ or Kilosort 2.0^52^ and manually curated. Only units that had an absolute refractory period of at least 1 ms were included in subsequent analysis. Behavioural variables were extracted from video recordings using DeepLabCut (DLC^53^; training set 1000 frames, 1 million training iterations). All timepoints during which the mouse was either inside the shelter or leaning down the behavioural platform where excluded from further analysis. Head-direction and shelter direction were calculated using DLC tracked ear-positions: head-direction was defined as the angle between the direction perpendicular to a line that joins both ears and the horizontal axis; shelter direction was defined as the angle between the same direction as above and a line connecting the centre point between the ears and the shelter entrance (Fig. 1A, Extended Data Fig. 2B, Supplementary Video 3).

#### Single cell tuning analysis: criteria for classification of shelter-direction and head-direction cells

We used Rayleigh vector length^54^ in head-direction and shelter direction spaces, as previously described in head-direction tuning testing in rodents^55^, to quantify the tuning of each cell to each of these variables. A neuron was considered to be significantly tuned to a variable when the length of its Rayleigh vector exceeded the 95th percentile of the distribution of Rayleigh vectors lengths computed for 1000 shuffled-datasets, generated by shifting recorded spike trains from all cells coherently by a uniform amount chosen randomly between 1 and 100 s. To ensure the tuning estimation was not influenced by potential behavioural biases, we binned the variable space into 16 bins and sampled them equally. Single units from RSP or SC were classified as shelter-direction cells if all the following criteria were met: i) for each epoch the neuron has to display a significant shelter direction Rayleigh vector; ii) for each epoch the neuron tuning to shelter direction must de decoupled from tuning to head-direction (using the Tuning Entanglement Decoupling (TunED) analysis, see below); iii) shelter direction tuning has to be stable across all epochs (above chance ROC decoding^56^ along the direction in which Rayleigh vector points during the shelter position 1 epoch, in shelter position 2 epoch); iv) shelter direction tuning must rotate with the shelter (after rotation of the shelter, the tuning to shelter position 2 must not be significantly explained by the tuning to shelter position 1, using TunED analysis). Single units from RSP or SC were classified as head-direction cells if all the following criteria were met: i) for each epoch (including the no shelter epoch) the neuron had to display a significant head-direction Rayleigh vector; ii) for each epoch the neuron tuning to head-direction had to be decoupled from tuning to shelter direction (using the TunED method); iii) head-direction tuning has to be stable across all epochs (above chance ROC decoding along the direction in which Rayleigh vector points during the first epoch of the experiment, in the subsequent periods).

#### Tuning entanglement decoupling (TunED) analysis

To determine whether a neuron that is statistically tuned to two correlated variables is driven by just one of the variables (*v*_*1*_, ‘driver’) while the tuning to the second variable (*v*_*2*_, ‘passenger’) is an artefact of the correlation, we performed tuning entanglement decoupling analysis (TunED). The method takes a list of spike-counts *r* along with simultaneously measured stimulus variables v_1_ and v_2_ Let

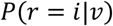

be the probability that the neuron’s spike count *r* takes a given value, given a discrete value for the variable *v*, the tuning curve of the neuron to stimulus variable *v* (observed tuning to *v*_*1*_ or *v*_*2*_, Extended Data Fig. 2A, right panel, dark yellow and dark blue) is then the conditional mean spike count

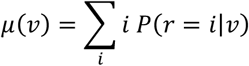

To determine whether a neuron is only tuned to the driver variable v_1_ or indeed significantly tuned to both *v*_*1*_ and *v*_*2*_, the method formulates the null hypothesis that the neuron’s activity is purely driven by the *v*_*1*_. In this case it can be shown that the tuning curve to *v*_*2*_ (*μ(v*_*2*_); expected tuning of *v*_*2*_ given tuning to *v*_*1*_, Extended Data Fig. 2A, right panel, light yellow and light blue) can be expressed as:

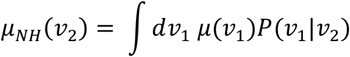

Where the suffix *NH* denotes this is not the actual tuning curve to *v*_*2*_ but instead it is derived by the above null hypothesis.

TunED analysis was used to classify cells has head direction or shelter-direction cells. Recorded spike and head direction and shelter direction time series were bootstrapped 1000 times, generating 1000 observed tuning curves to each variable (Extended Data Fig. 2C, left panels, dark yellow and dark blue) as well as 1000 expected tuning curves of the same variable using the null-hypothesis equation above (Extended Data Fig. 2C, left panels, light yellow and light blue). For each variable, the Euclidean distance (d_HSA_ and d_HD_) between the observed (OT_HSA_ and OT_HD_) and the expected (ET_HSA_ and ET_HD_) tuning curves was calculated for each bootstrap iteration (Extended Data Fig.2C, right panels, yellow and blue histograms). Finally, we computed the distribution of the differences between each variable d_HSA_ and d_HD_ for each bootstrap iteration (Extended Data Fig. 2C, right panels, dark grey histograms). If the distribution of d_HSA_ - d_HD_ was significantly smaller than zero (both 2.5^th^ and 97.5^th^ percentile <0, vertical dotted lines in Extended Data Fig. 2C, right panels) the cell was considered a head direction cell (Extended Data Fig. 2C, top panels); if the distribution of d_HSA_ - d_HD_ was significantly larger than zero (both 2.5^th^ and 97.5^th^ percentile > 0) the cell was considered a shelter direction cell (Extended Data Fig. 2C, bottom panels); otherwise the cell was not considered a shelter direction nor head direction cell.

#### Generalized linear model (GLM)

Generalized linear models^57^ were used to predict the probability of spiking at certain time *t* during shelter position 1 and shelter position 2 epochs, given the values of a set of simultaneously recorded variables at time *t* (head-direction, shelter direction, locomotion velocity, head angular velocity, change in head direction within the next 100ms and 200ms; see ^22^). For each of these variables we defined a set of equally spaced bins (locomotion speed, 11; head direction, 27; shelter direction, 27; head angular velocity, 23; change in head direction within the next 100ms, 13; change in head direction within the next 200ms, 13). A time varying binary predictor was then defined for each bin (excluding one) of each variable. At a given *t*, each binary predictor could be 1, if the value of the respective behavioural variable fell in that bin at *t* and zero otherwise. The GLM was fitted with the *glmfit* MATLAB function assuming logistic link function and Bernulli probability distribution. Model prediction accuracy was assessed by performing 10-fold cross-validation and computing Pearson correlation coefficient between predicted and real spike trains (smoothed over 100 ms). The analysis above was also performed after excluding shelter direction from the model.

#### Population decoding analysis

We employed multiclass linear discriminant analysis (LDA; ^58^) to decode shelter direction from spike trains of RSP and SC neural populations, and change in head direction within the next 100ms from spike trains of SC neural population. Shelter direction was binned into 16 equally populated classes (bin range: −180° to 180°, bin amplitude: 22.5°), and change in head direction into 9 classes (bin range: −27° to 27°, bin amplitude: 6°). Population data was constructed by grouping all recorded neurons and aligning their activity according to the value of shelter direction (or change in head direction) in which it was measured. Both shelter positions 1 and shelter positions 2 epochs were divided into 6 interleaved time periods of equal duration; for each epoch periods 1, 3 and 5 were used to train the classifier, while periods 2, 4 and 6 were used to test its accuracy. Prediction accuracy was computed as the fraction of observations of the testing set classified in the main diagonal of the classifier confusion matrix and in the adjacent ones (± 1), over the total number of observations in the testing set. Predictions accuracy was also compared to the one obtained for a shuffled dataset.

### Single unit recordings paired with chemogenetic loss-of-function analysis

#### Firing rate change

The effect of inactivation was assessed separately in RSP and SC neurons. Change in firing rate index (Δ firing rate index) for each neuron in the period before (shelter position 2 epoch) and after the administration of CNO (or saline) was computed as:

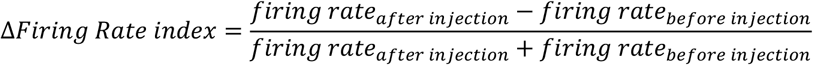

#### Single cell analysis

To test for changes in the percentage of neurons classified as shelter-direction neurons we first identified shelter-direction as described above in the periods before and after the injection.

We then computed:

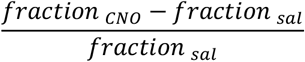

where *fraction* is the fraction of shelter-direction cells after the injection.

#### Population analysis

We computed the change in prediction accuracy of multiclass LDA classifiers (see *population decoding analysis*), following injection of CNO or saline in respect to baseline. The period before injection (including shelter positions 1 and 2 epochs) was divided into 12 bins of equal duration. Odd bins were used to train the classifier and even bins used to cross-validate it, producing the baseline prediction accuracy result. The same classifier was used to predict the behavioural variable, given spike trains, after injection of CNO or saline. This was repeated 10 times, by randomly choosing the samples of data for each class. To probe the effect of inactivation of RSP, we tested the statistical significance of the difference between the change the classifier’s prediction accuracy after CNO and after saline injection, using *sign rank test*. The first 35 min after injection were discarded from these analyses. Single unit recordings paired with chemogenetic loss-of-function experiments were performed with two mice (see Laurens 2022^59^). The statistical power supporting the conclusion was reached because of the high number of recorded neurons (see *Results*).

### Single unit recordings in head fixed mice paired with dual opsin-assisted circuit mapping and opto-tagging

Neuropixels silicon probes (phase3A, option3, 384 channels) were used to record extracellular spikes from SC neurons of 3 adult VGluT2::cre and 4 adult VGAT::cre mice, previously injected (6-8 weeks) as described in *Experimental procedure: Viruses*. A metal custom-made head-plate and optic fibre (Newdoon, see Supplementary table 3) were attached to the skull using dental cement. A craniotomy was made over the SC and sealed with Kwik-Cast on the day of the first recording session. During each recording session, mice were placed on a plastic wheel and head fixed. We performed up to four recording sessions per mouse. Before insertion, the probe was coated with one of the four following dyes, to identify and distinguish probe tracks: DiI (1 mM in ethanol, Invitrogen), DiO (1 mM in ethanol, Invitrogen), Vybrant™ DiD (1 mM in ethanol, Invitrogen), Cellbrite blue (Cytoplasmic Membrane-Labeling Kit - Biotium). Craniotomy and skull surface were submerged with aCSF (see *Experimental procedures: In vitro whole cell recordings*) and a coated (*AgCl*) silver wire (0.35 mm diameter, Goodfellow) was held in the aCSF bath for external referencing and grounding. For recording, the probe was slowly inserted into the SC (see Fig. 4A) to a depth of 2.8–3.0 mm and left in place for at least 45 min before the beginning of the recording session. Data was acquired as described in *Experimental procedure: Single unit recordings in freely moving animals* section.

Dual opsin-assisted circuit mapping and optotagging was performed analogously to what described in *Experimental procedures: In vitro whole cell recordings* section using a Vortran Stradus VersaLase as light source (wavelengths: 472 nm and 639 nm), except that for ChR activation light pulse was 5 ms long (power at fibre tip ∼10mW) and for ChrimsonR activation we used a 20 Hz train of 10 ms – long pulses (1 sec duration, power at fibre tip ∼10mW). These stimuli were repeated interleaved 50 times with an interstimulus interval of 35 seconds. A neuron was classified as putative vGlut2^+^ or a vGAT^+^ if it spiked in at least 75% of the 472nm pulses. Population dynamics shown in Fig. 4B were robust to the additional constraints of the first spike latency having small jitter across trials (IQR 1ms and 2.5ms).

### Single unit recordings in freely moving mice paired with dual opsin-assisted circuit mapping and opto-tagging

Recordings and analysis in freely moving mice were performed as described in of *Experimental procedure: single unit recording in freely moving mice* in 2 adult VGluT2::cre and 2 adult VGAT::cre mice, previously injected (6-8 weeks) as described in *Experimental procedure: Viruses*. An optic fibre (Newdoon, see Supplementary Table 3) were attached to the skull on the same day of the Neuropixels chronic implant. At the end of each recording session, dual opsin-assisted circuit mapping and opto-tagging was performed as describe in *Experimental procedure*: *Single unit recordings in head fixed mice paired with dual opsin-assisted circuit mapping and opto-tagging*.

### Spiking Recurrent Neural Network modelling

An artificial spiking neural network with biologically constrained synaptic properties and connectivity was used to model the shelter direction RSP to SC circuit neural dynamics. The model was implemented in custom Python code and used the Brian2 neural network simulator package^60^. The simulated network was composed of three distinct populations (representing RSP, SC excitatory and SC inhibitory units) of 512 conductance-based leaky integrate and fire (LIF) neurons adapted from previous models of spiking neural networks with ring-attractor dynamics^61^. The neuronal network model was designed to implement a ring like-neural network, with neurons in each population numbered (1-512) and ordered such that nearby neurons corresponded to similar angles in the ring^61^. A *connectivity kernel* was used to determine which neurons in the source population projected to which neurons in the target one and the strength of the connection between each pair (*w*), based on their position in the ring. Two kernel types were used. *Diagonal* created connections with a kernel shaped as a Gaussian distribution (of standard deviation equal to *kernel width* – see Supplementary Table 4), such that entries in the main diagonal had the strongest connection, the first super- and sub-diagonals weaker connections etc with weights approaching zeros for entries further away from the main diagonal. *Inverted diagonal* was analogous to the *diagonal* except that synaptic weight was highest for off-diagonal elements and approached zero for elements on the main diagonal with a Gaussian kernel^27^. Within the constraints of the connectivity kernel, synaptic connections between neurons in the three populations were randomly created with a probability *p*_*connection*_(see below).

The membrane potential (*V*_*m*_) dynamics of each neuron was modelled by the following ODE:

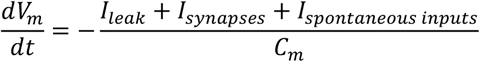

where *I*_*leak*_=*g*_*leak*_∗(*V*_*m*_− *E*_*leak*_) is the leakage current, *I*_*spontaneous inputs*_ is the spontaneous synaptic current and *I*_*synapses*_ is the sum of synaptic currents (*I*_*synapse*_) from each presynaptic population: RSP inputs (*I*_*rsp*_), excitatory SC inputs (*I*_*vglut sc*_) and inhibitory SC inputs (*I*_*vgat sc*_).

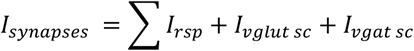

In order to better fit experimental data, each *I*_*synapse*_ was modelled with a slow (*I*_*slow*_) and a fast (*I*_*fast*_) component:

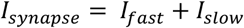

For each component, the dynamics in synaptic conductance (g) is governed by the time constants *τ*_*rise*_ and *τ*_*decay*_

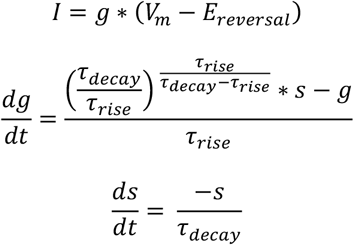

Following a presynaptic spike, the value of *s* at the synapse between the presynaptic neuron *i* and the postsynaptic neuron *j* is incremented by *g*_*max*_(*i, j*).

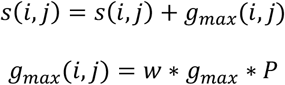

such that g at any time t after a presynaptic spike is given by:

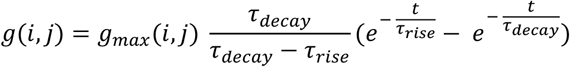

*P* models the short term plasticity dynamics (*P = u* ∗*x*), where *x* is the proportion of available synaptic resources at time of presynaptic spike (initialised to 1) and *u* the release probability, following the equations described in Tsodyks and Markram 1997^62^. *P* was set to 1 for the slow component, such that only the fast component shows short-term plasticity. The following variables ^63^obtained from either our own experimental data (directly or after compartmental modelling in NEURON^63^) or from published data ^3,64^ where necessary: synaptic properties - maximum connection probability of the synapse type, *g*_*max*_, *τ*_*rise*_ and *τ*_*decay*_ of the fast and slow components; ^3,64^ cellular properties: membrane resistance (*g*_*leak*_ ^*-1*^) and capacitance (*C*_*m*_), resting membrane potential, and spike threshold and spike reset potential.

To mimic the effect of optogenetic stimulation of RSP neurons on neural dynamics in SC, a randomly selected subset of RSP neurons received an external input in the form of a train of positive current. Stimulation patterns were the same as the ones described in *Experimental procedures: Single unit recordings in head fixed mice paired with dual opsin-assisted circuit mapping and opto-tagging*. An exhaustive grid search was used to estimate parameters yielding neural dynamics similar to those measured experimentally. Briefly, for each combination of parameters a neural network model was simulated as above and the normalised average activity of vGluT2^+^ and vGAT^+^ neurons was measured and compared to the experimentally recorded neurons calculating the sum of squared error. To generate shelter direction tuning curves, 10 non-overlapping subgroups of nearby RSP neurons were defined, each spanning a 36 degrees angle. Each RSP subgroup was stimulated individually by increasing the strength of the spontaneous inputs to its neurons for 1 second, and the simulated spike trains of SC vGAT^+^ and vGluT2^+^ neurons were used to compute tuning curves and Rayleigh vectors as described in *Experimental procedure: Single unit recordings in freely moving animals*.

### General data analysis

Data analysis was performed using custom-written routines in Python 2.7 and MATLAB and custom code will be made available on request. Data are reported as median ± i.q.r. or mean ± s.e.m unless otherwise indicated. Statistical comparisons using the significance tests stated in the main text were made in GraphPad Prism, MATLAB, R and statistical significance was considered when *P* < 0.05. Data were tested for normality with the Shapiro–Wilk test, and a parametric test used if the data were normally distributed, and a non-parametric otherwise, as detailed in the text next to each comparison. To test whether a population of angular measures was distributed non-uniformly around the circle we used a V-test for non-uniformity of circular data. The z-test for equality of proportion was used to compare proportions. Unless otherwise stated, statistical difference between behavioural metrics in CNO and control experiments was computed using the following permutation test (developed in collaboration with J. Rapela, Gatsby Data Center). Two groups of mice M and N, each of numerosity *m* and *n*, were tested under a different experimental condition (e.g.: control or CNO). The test statistic computed is the difference between the mean of all trials pooled in CNO group and the mean of the trials in the control group. For the permutation, at every iteration each mouse is randomly re-assigned to CNO or control group, while keeping the total number of mice per condition the same. The N number in this statistical comparison is therefore the number of mice in each group. The procedure was repeated 100000 times and the test statistics was computed after each iteration, obtaining the null distribution of the test. The p-value of the test is one minus the percentile of the null distribution corresponding to the value of the test statistic of the non-shuffled data.

The data and code that support the findings of this study are available from the corresponding authors upon request.

## Acknowledgments

This work was funded by a Wellcome Senior Research Fellowship (214352/Z/18/Z) and by the Sainsbury Wellcome Centre Core Grant from the Gatsby Charitable Foundation and Wellcome (090843/F/09/Z) (T.B.), MRC PhD Studentship (R.V.), Boehringer Ingelheim Fonds PhD fellowship (R.V.), Gatsby Unit/SWC Joint Research Fellowship in Neuroscience (D.C.), UCL Wellcome 4-year PhD Programme in Neuroscience (O.P.A), A*STAR National Science Scholarship (PhD; Y.L.T) and the SWC PhD Programme (Y.L.T.). We thank members of the Branco lab and T. Mrsic-Flogel for discussions; J. Rapela and P. Shamash for advice on statistical analysis; A. Murray, I. Duguid, C. Schimdt-Hieber, N. Burgess, M. Tripodi, and the I. Bianco lab for comments on the manuscript; M. Strom, T. Okbinoglu, R. Campbel, the SWC Neurobiological Research Facility and FabLabs for technical support; K. Betsios for programming the data acquisition software; T. R. Stones for inspiration; G. T. Gray for viral constructs. Source of mouse silhouettes: https://scidraw.io/.

## Author contributions

D. C and R.V. performed behavioural experiments. D.C., F.C. and Y. L. T.: theoretical modelling; D.C.: single-unit recordings; O.P., A. V. S. and Y. L.T.: in vitro electrophysiology; D.C., P. I., S. K., R. V.: surgeries. D.C., O.P., R. S. P., Y. L.T and R. V. data analysis; D. C., P. I., Y. L. T. and R. V.: histological preparations and imaging; T.B., D. C., Y. L. T. and R.V. experimental design; T.B. and R.V. conceived the project. T. B. wrote the manuscript, with critical input from D. C., T. W. M, Y. L. T and R. V.

