## Supplementary Figures for "A cortico-collicular circuit for accurate orientation to shelter during escape"

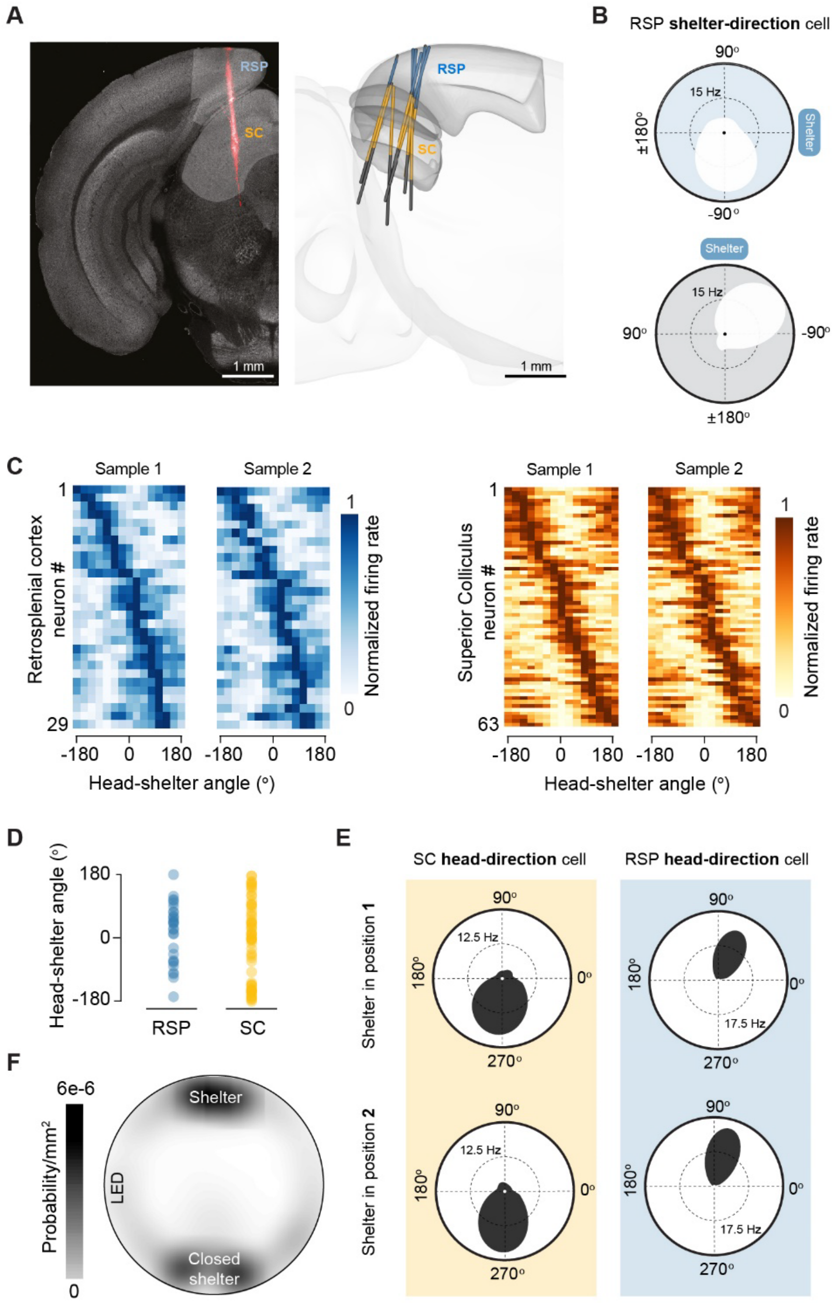


**Extended Data Figure 1 - Single unit recordings of shelter-direction and head-direction cells**

(**A**) Left: Coronal image of post-recording histology showing the track of the neuropixels probe (red). Right: 3D rendering of probe tracks in all chronically implanted mice (**B**) Example tuning curves for a shelter-direction neuron in the RSP before and after shelter rotation. (**C**) Tuning curves for non-overlapping subsets of data generated by random sampling. For RSP and SC, the plot on the left is sorted by tuning peak, and the sorting indexes have been used to sort the plot on the right. (**D**) Summary plot of preferred tuning angle for RSP and SC shelter-direction neurons. (**E**) Example tuning curves for allocentric head-direction neurons in the SC and RSP. We recorded 2% head-direction cells in the SC, and 6% in the RSP. **(F)** Summary plot showing the occupancy of the arena during exploratory behaviour in the presence of a second, closed shelter.

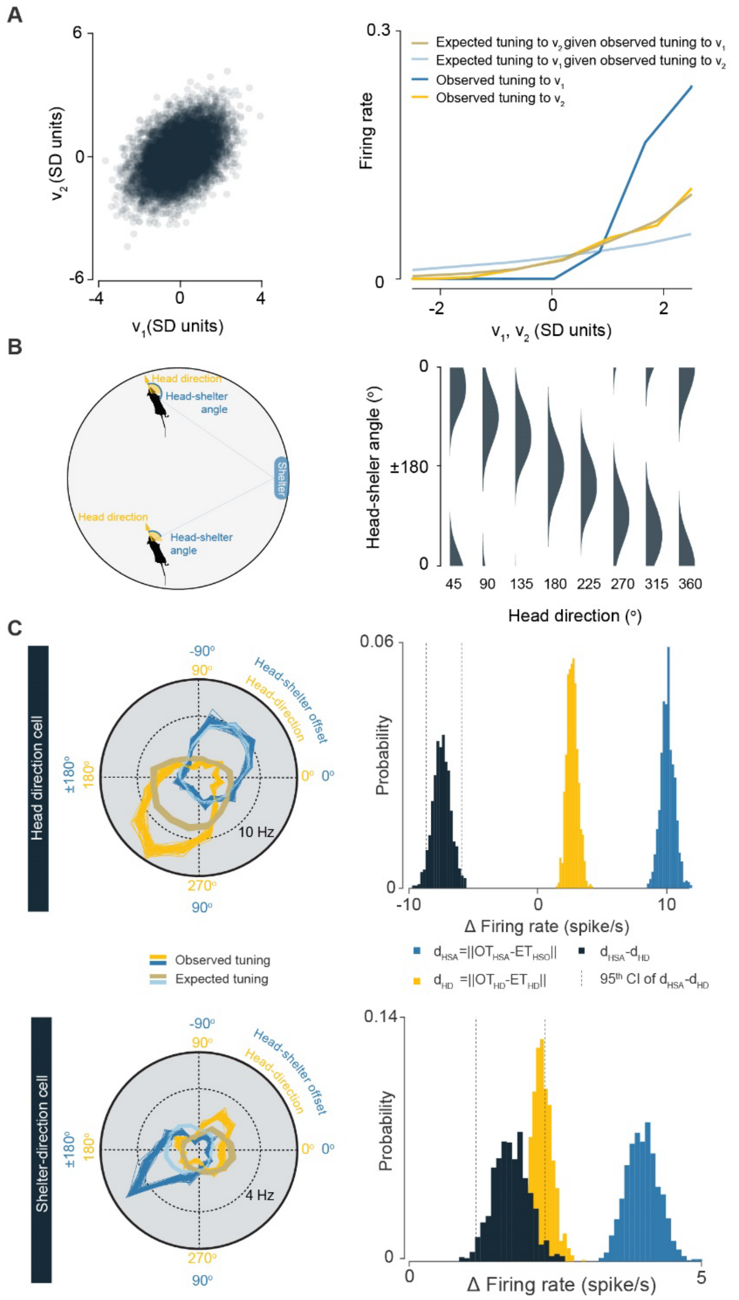


**Extended Data Figure 2 - Tuning entanglement decoupling analysis**

(**A**) Left, plot illustrating a driver variable (v1) and a correlated passenger variable (v2; Pearson's correlation coefficient = 0.45). v1 samples are drawn from a normal distribution of mean 0 and standard deviation 1. The jth sample of v2 is computed as v_2j=0.5* v_1j+ε_j, where each ε_j is drawn from a second normal distribution with mean 0 and standard deviation 1. Right, v1 is used to simulate the spiking of a neuron such that the probability of firing is equal to 0.1 * v1 if v1>1 and 0 if v1≤1. TunED analysis was then applied to v1, v2 and simulated spiking data as described in Methods to compute observed and expected tuning curves to v1 and v2. The method correctly identifies v1 as the driver variable. The observed tuning curve to the passenger variable v2 (dark yellow) can be fully explained by the tuning to the driver variable (brown). In contrast, the observed tuning to driver variable (dark blue), cannot be explained by the tuning to the passenger variable (light blue). (**B**) Left, schematic of head-shelter angle and head direction variables during the experiment. Right, correlation between head-shelter angle and head direction in our experimental settin plotted for eight values of head direction for each grid location. (**C**) Left, tuning curves of neurons for which the driver variable was head direction (top) or head-shelter angle (bottom). Right, illustration of the statistical method used to determine whether the driver variable of the neuron was head shelter offset, head direction or none of them (see Methods for details). Briefly, the distribution of dHSA - dHD (dark grey histogram) indicates whether the expected and observed tuning curves are more similar for head-shelter angle or for head direction. If the dHSA - dHD distribution is significantly smaller than zero (both 2.5th and 97.5th percentile <0, vertical dotted lines) the cell is classified as a head direction cell; if the distribution of dHSA - dHD is significantly larger than zero (both 2.5th and 97.5th percentile > 0) the cell is classified as a head-shelter angle cell; otherwise the cell is not considered a shelter-direction nor head direction cell.

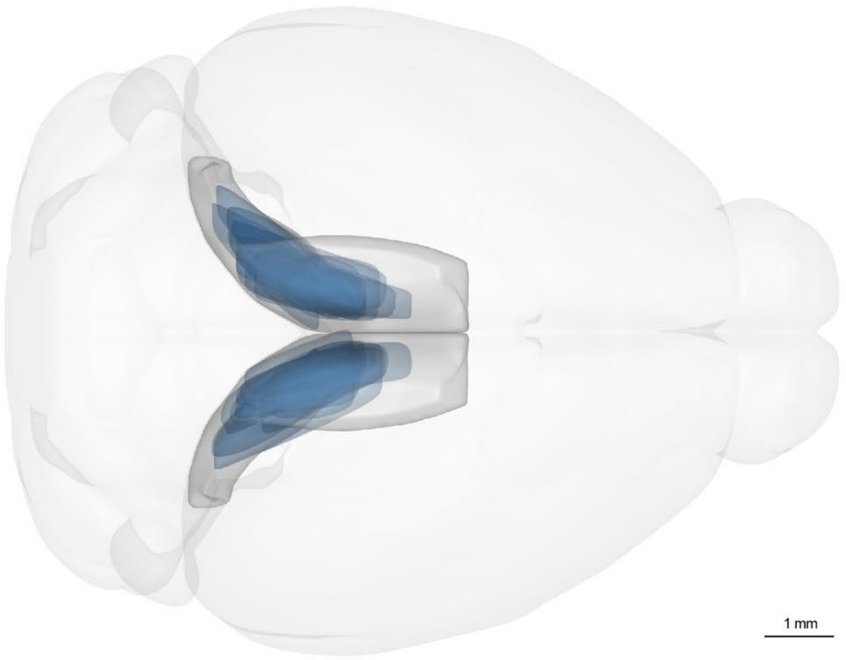


**Extended Data Figure 3 - Histology for loss-of-function of SC-projecting RSP neurons**

3D rendering of the location of SC-projecting RSP neurons expressing hM4Di for the mice in the following datasets: escape behaviour assay, orientation to sound assay, food-seeking assay, single unit recondings during chemogenetic inactivation. For an example coronal section see Fig. 2A.

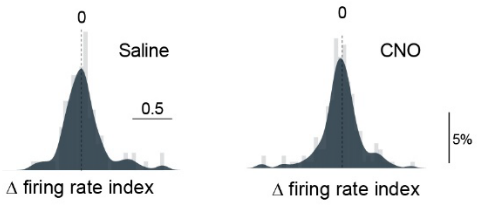


**Extended Data Figure 4 - RSP loss-of-function does not affect average SC firing rates**

(**A**) Population histograms for firing rate of SC single units after saline and CNO i.p. injection in animals expressing hM4Di in SC-projecting RSP neurons P=0.75 one-tailed Kolmogorov-Smirnov test; N=264 units, 2 mice;

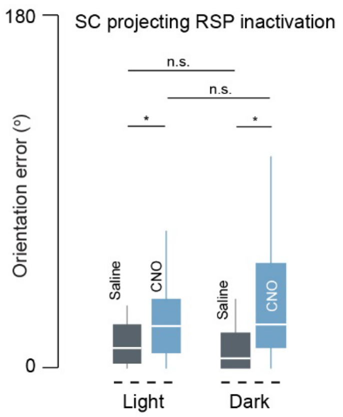


**Extended Data Figure 5 - Shelter orientation error does not depend on the enviroment luminance level**

Shelter orientation error increses both during light and dark conditions upon inactivation of SC-projecting RSP neurons, in comparison to saline control (dark: permutation test p-value 0.0302; saline n = 5 mice 24 trials; CNO 11 mice 58 trials. light: permutation test p-value 0.038; saline n = 6 mice 27 trials; CNO 9 mice 47 trials). No significant differences were observed between saline in light and dark condition (permutation test p-value 0.46) or CNO in light and dark condition (permutation test p-value 0.26).

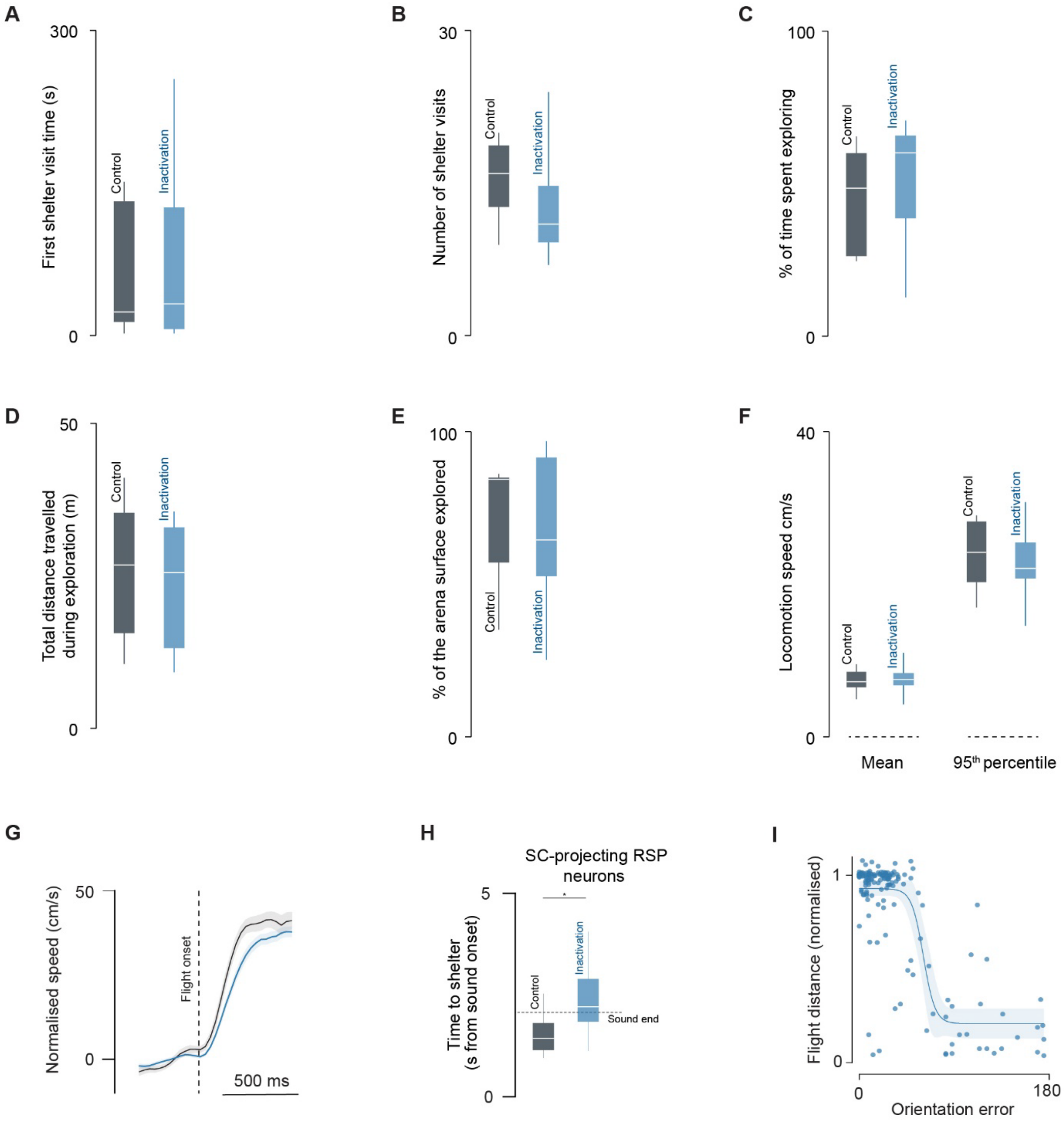


**Extended Data Figure 6 - Additional analysis of the effect of RSP-SC loss-of-function on behaviour**

(**A-F**) Navigation during exploratory behaviour is not affected by inactivation of SC-projecting RSP neurons. Panels show quantification of exploratory behaviour during the time period preceeding the presentation of the first threatening stimulus for saline control (black, N=6) and CNO (blue, N=11) mice, expressing hM4Di in SC-projecting RSP neurons. None of the metrics differs between the two groups (2-tailed Mann-Whitney test, $p>0.15$ for all metrics). (**A**) Latency between the beginning of the experiment and the first time the mouse entered in the shelter. (**B**) Number of times the mouse entered in the shelter. (**C**) Percentage of time the mouse spent outside the shelter. (**D**) Total lenght of the path travelled while outside the shelter. (**E**) Percentage of the arena surface explored while outside the shelter. (**F**) Average and 95$^{th}$ percentile of mouse locomotion speed while outside the shelter. The duration of the time period preceeding the presentation of the first threatening stimulus did not differ between saline control and CNO groups (2 tailed Mann-Whitney test, $p = 0.51$). (**G**) Average change in speed after threat presentation for saline control (black) and CNO (blue) showing that both groups of mice initiate escape with similar vigour. (**H**) Summary data for time to reach the shelter after escape initiation. (**I**) Length of flight after escape initiation as a function of orientation error, showing that larger errors are associated with shorter flights. (Fitted function: Boltzmann sigmoidal equation; slope -5.5, $p=0.02$; F-statistic goodness of fit test against constant model, $p<0.0001$). Shaded area: 95% confidence interval. Distance is normalised to the distance to shetler at escape onset .

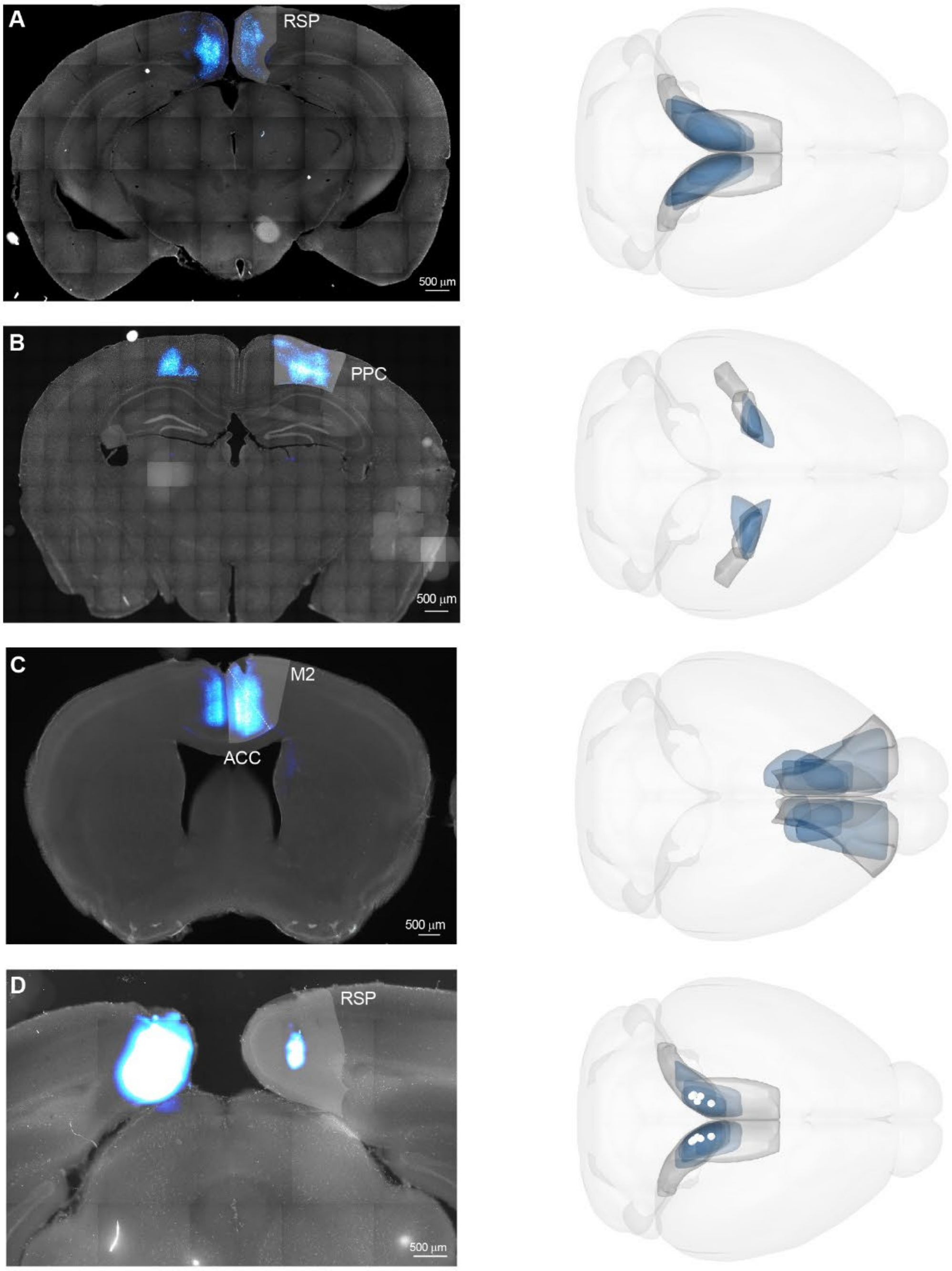


**Extended Data Figure 7 - Histology for cortical loss-of-function**

Coronal sections and 3D renderings showing neurons targeted with hM4Di expression in the entire RSP (**A**), posterior parietal cortex (**B**) and anterior motor areas (**C**). **D** shows fluorescently-labelled muscimol targeted to the RSP. White cylinders represent infusion cannulae location.

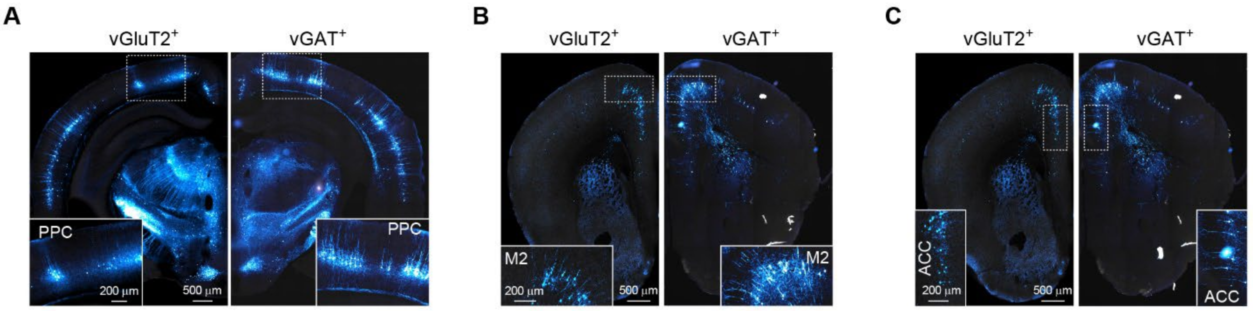


**Extended Data Figure 8 - Additional cortical inputs onto SC neurons**

Coronal images of monosynaptic rabies tracing from starter SC cells in excitatory (vGluT2$^+$) and inhibitory (vGAT$^+$) neuron populations, showing prominent inputs from the posterior parietal cortex (**A**), M2 (**B**) and anterior cingulate cortex (**C**).

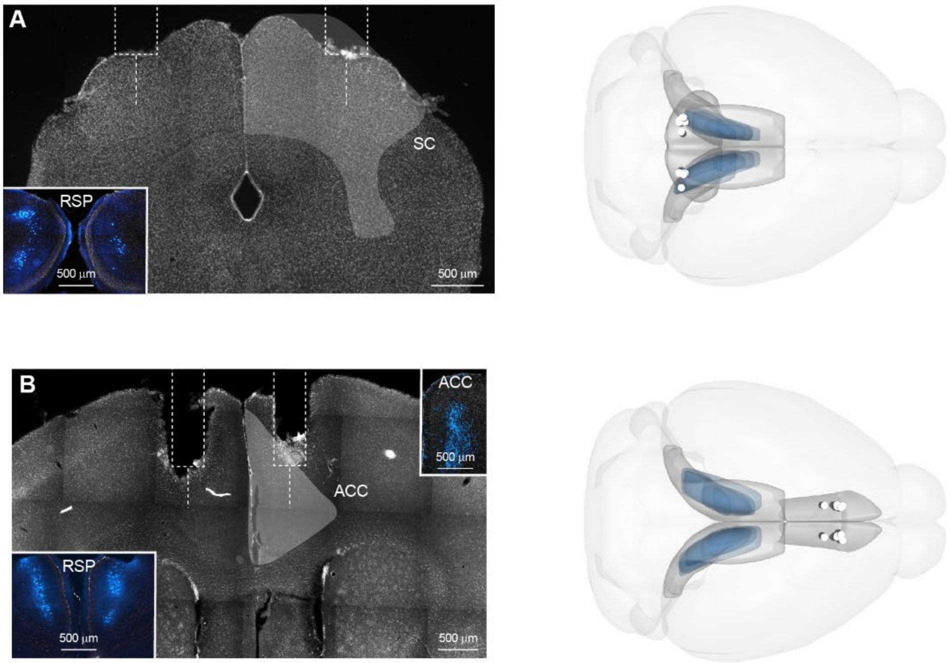


**Extended Data Figure 9 - Histology for projection-specific RSP-SC loss-of-function**

Coronal sections and 3D renderings showing guide, internal cannulae locations (white dotted lines and white cylinders) implanted in the superior colliculus (SC; **A**) and anterior cingulate cortex (ACC; **B**). Insets and blue shades in the right panels show SC-projecting RSP neurons of the respective mouse, expressing hM4Di. Inset in B (top-right) shows the axon collaterals to ACC of SC-projecting RSP neurons.

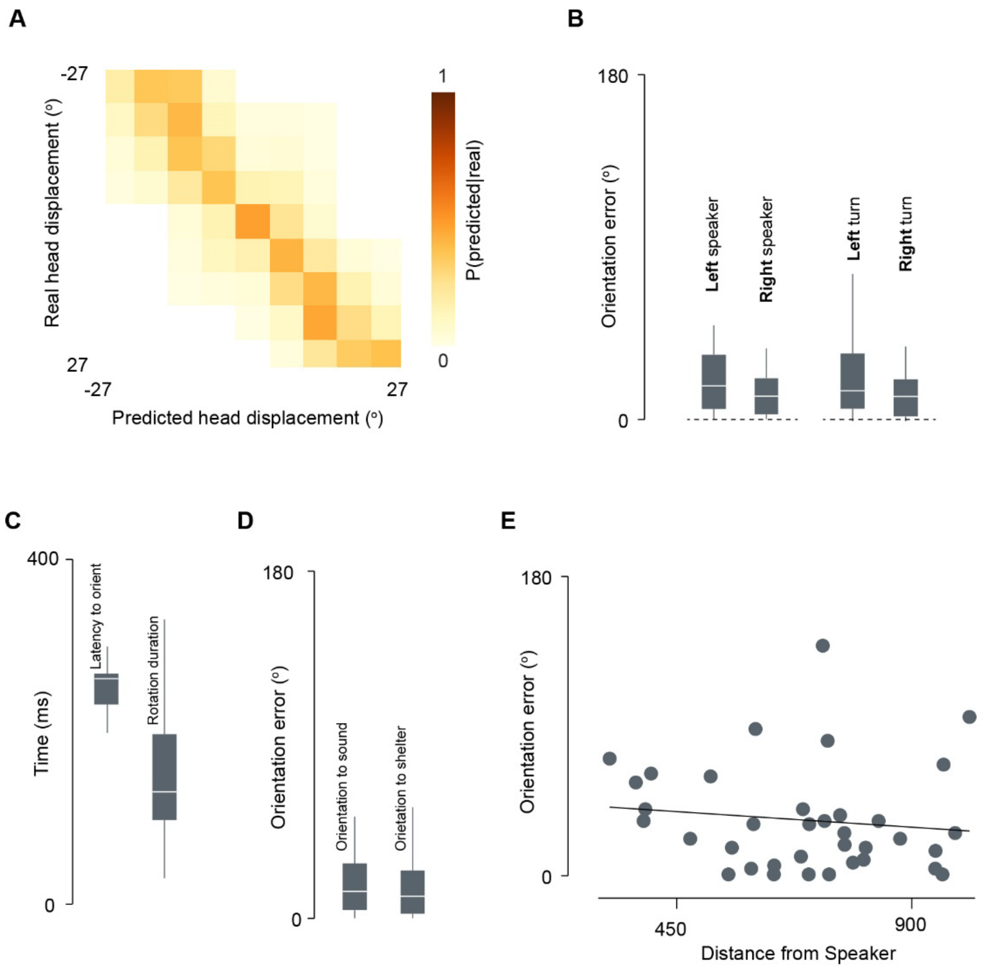


**Extended Data Figure 10 - Quantification of head-displacement prediction and orientation to sound performance**

(**A**) Cross validated confusion matrix for LDA population decoding of the angle of future head displacement (100 ms ahead) from SC firing rates (prediction accuracy 0.78). (**B**) Summary data showing that mice orient accurately to sound (with no biases for left or right speaker and for left or right turns; permutation test P = 0.27 and P = 0.33 respectively, N = 36 , 6 mice), with short latencies (**C**, left) and fast movements (**C**, right). (**D**) Summary data showing that mice are equally accurate when orienting to sound or to shelter (permutation test P = 0.82; orientation to sound N = 36, 6 mice; orientation to shelter N = 32, 5 mice). (**E**) Orientation to sound accuracy does not depend on the distance at sound onset between the mouse and the speaker (slope = -0.012, P = 0.48, linear regression).

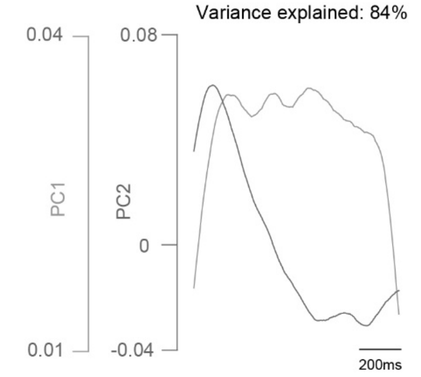


**Extended Data Figure 11 - Principal components of SC population dynamics during RSP activation**

Principal component 1 and 2 of SC vGAT$^+$ and SC vGlut2$^+$ neurons during cortical activation (same data as Figure 4B). The first two principal component are sufficient to explain most of the variance present in the data (84%) and closely resemble the temporal dynamics observed for SC vGAT$^+$and SC VGlut2$^+$ neurons.

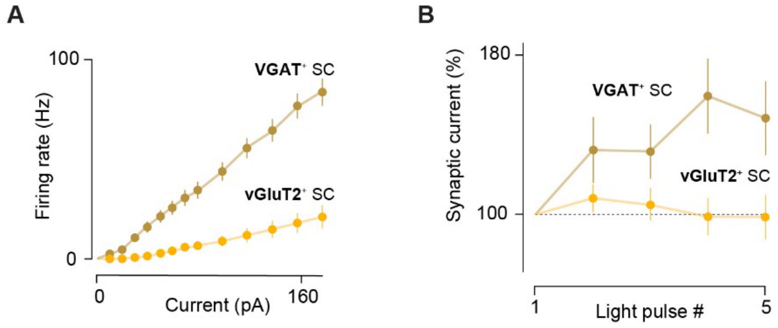


**Extended Data Figure 12- Biophysical properties of SC neurons receiving RSP input**

(**A**) Summary curves for action potential firing from somatic current injection. (**B**) Summary data for short-term plasticity of RSP inputs onto SC neurons

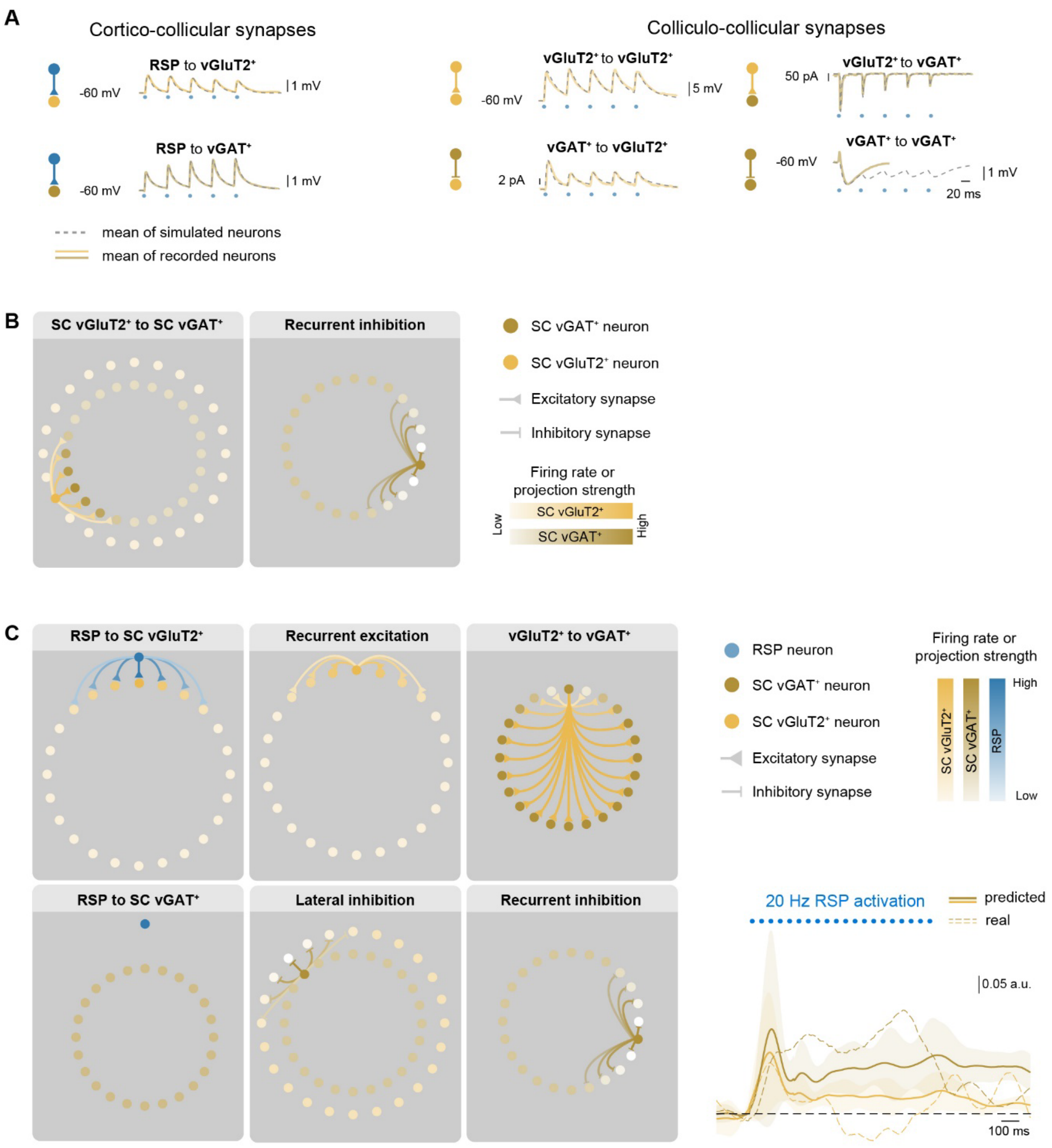


**Extended Data Figure 13 - Elements of the lateral feedforward inhibition and ring attractor models**

**(A)** Real and simulated synaptic currents or potentials for all synaptic connections in the model. **(B)** Additional circuit elements of the feedforward lateral inhibition model (c.f. Figure 5D). **(C)** Left: Circuits element of the standard ring attractor model. Right: predicted firing rate of vGluT2⁺ and vGAT⁺ SC populations following 20 Hz activation of RSP neurons in the model, compared to observed dynamics (dashed lines, see also in Figure 4B).

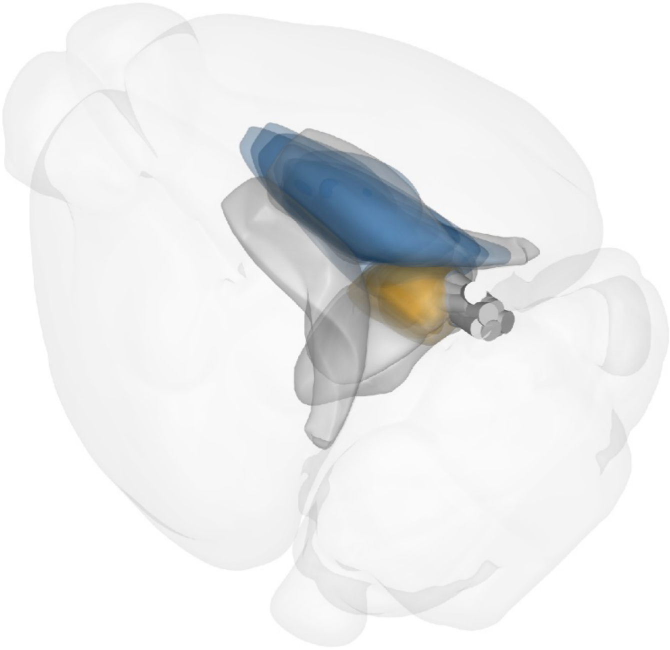

**Extended Data Figure 14 - 3D reconstruction of viral injection and fiber placement of dual opsin-assisted circuit mapping and optotagging**

3D reconstructions of the location of Chrimson expressing neurons in the entire RSP (blue), ChR2-expressing SC vGlut2$^{+}$ and vGAT$^{+}$ neurons (yellow), and optic fibers (white cylinders) used in the freely moving and head-fixed dual-opsin and optotagging experiments. For an example coronal section see Figure 4A.
