## Supplementary Tables for "A cortico-collicular circuit for accurate orientation to shelter during escape"

| Target region | Coordinates(mm) | | | Cannula (mm) | | Volume(μL) |
|---|---|---|---|---|---|---|
| | ML | AP | DV | Dual | Internal | |
| Superior colliculus | ± 1.00 | - 0.30λ | -1.20 | 2.00 | -0.50 | 0.8 – 1.2 |
| Anterior Cingulate Cortex | ± 0.60 | +1.10b | -1.10 | 1.20 | -0.50 | 0.8 – 1.2 |
| Retrosplenial Cortex | ± 0.60 | +0.4 λ | -0.8 | 1.20 | -0.50 | 0.8 – 1.2 |

**Supplementary Table 1 - Parameters for cannula implantation and drug delivery in pharmacological and chemogenetic inactivation experiments.** Cannula implant coordinates are specified from bregma (b) or lambda (λ). The final target depth is the sum of the DV and internal lengths. Dual indicates the width between cannulae.

| Target region | Coordinates(mm) ML | AP | DV | Volume(nl) |
|---|---|---|---|---|
| Superior Colliculus | ± 1.10 | - 0.40λ | -1.90 | 100-120 |
| | ± 1.00 | -0.40λ | -1.50 | 100-120 |
| | ± 1.00 | 0.00λ | -2.00 | 100-120 |
| Retrosplenial Cortex | ± 0.65 | +0.40λ | -1.00; -0.70 | 100-120[1] |
| | ± 0.40 | +1.10λ | -1.20; -1.90 | 100-120[1] |
| Anterior Motor Areas | ± 0.70[2] | +0.26b[2] | -1.05[2] | 180[2] |
| | ± 0.75[2] | +1.34b[2] | -1.20[2] | 180[2] |
| | ± 0.70[3] | +1.10b[3] | -1.40[3] | 180[3] |
| | ± 0.85[4] | +1.41b[4] | -1.60; -1.10[4] | 180[4] |
| | ± 1.10[4] | +1.93b[4] | -1.65; -1.10[4] | 100-120[4] |
| | ± 1.80[4] | +2.52b[4] | -2.00; -1.50[4] | 100-120[4] |
| | ± 1.00[4] | +2.52b[4] | -1.75; -1.25[4] | 100-120[4] |
| | ± 0.25[5] | +1.10b[5] | -1.80[5] | 180[5] |
| Posterior Parietal Cortex | ± 1.25 | +1.88λ | -0.70 | 100-120 |
| | ± 1.7 | +1.88λ | -0.70 | 100-120 |

[1]For Chr2-assisted circuit mapping the volume of AAV1-CAG-hChR2(H134R)-mCherry-WPRE-SV40 injected was 60-75nl.

[2-5] Combinations of injection coordinates used in different mice.

**Supplementary Table 2 - Parameters for viral injections experiments.** Injection site coordinates are specified from bregma (b) or lambda (λ). Parameters for retrograde rabies tracing have been specified in the main text.

| | Coordinates | | | | Optic fiber | |
|---|---|---|---|---|---|---|
| Target region | ML (mm) | AP (mm) | DV (mm) | Angle (°) | Diameter (μm) | NA |
| Superior colliculus (Insertion point) | + 0.75 | - 1.69λ | 0 | 40 | 400 | 0.5 |
| Superior colliculus (Fiber tip) | + 0.75 | -0.19 λ | -1.67 | 40 | 400 | 0.5 |

**Supplementary Table 3 - Parameters for optic fiber implantation for dual opsin-assisted circuit mapping and optotagging in head-fixed and freely moving mice.** AP coordinates are specified from lambda (λ).

| Cellular properties | RSP | SC $vGlut2^+$ | SC $vGAT^+$ |
|---|---|---|---|
| Spike threshold (*mV*) | -50 | -37.665 | -36.469 |
| Refractory period (*ms*) | 2 | 2 | 2 |
| Capacitance (*nF*) | 0.5 | 0.0468 | 0.03922 |
| Membrane resistance (GΩ) | 0.04 | 0.056 | 0.375 |
| Resting membrane potential (*mV*) | -70 | -60 | -60 |
| Reset potential (*mV*) | -60 | -62.341 | -60.664 |

| Synaptic properties | RSP to $vGAT^+$ | RSP to $vGlut2^+$ | $vGAT^+$ to $vGAT^+$ | $vGAT^+$ to $vGlut2^+$ | $vGlut2^+$ to $vGAT^+$ | $vGlut2^+$ to $vGlut2^+$ |
|---|---|---|---|---|---|---|
| $g_{max; fast}$ (*nS*) | 0.47 | 3.9 | 1.69 | 14.68 | 1.12 | 28.7 |
| $g_{max; slow}$ (*nS*) | 0.001 | 0.257 | 0.059 | 2.1 | 0.0609 | 0.504 |
| $\tau_{rise; fast}$ (*ms*) | 0.689 | 2.413 | 2.0 | 5 | 0.8 | 1 |
| $\tau_{rise; slow}$ (*ms*) | 33.0077 | 9.771 | 40 | 29.997 | 2.7997 | 45.3585 |
| $\tau_{decay; fast}$ (*ms*) | 0.718 | 3.384 | 19.5 | 6 | 1.3 | 19 |
| $\tau_{decay; slow}$ (*ms*) | 33.011 | 9.772 | 45.376 | 30 | 2.8 | 45.363 |
| **Synaptic properties: short term plasticity** | | | | | | |
| $\tau_{depression}$ (*ms*) | 40 | 700 | 120 | 120 | 330 | 120 |
| $\tau_{facilitation}$ (*ms*) | 160 | 25 | 1 | 1 | 25 | 50 |
| *u(0)* | 0.1 | 0.1 | 0.65 | 0.65 | 0.3 | 0.15 |
| **Synaptic properties: connectivity** | | | | | | |
| Kernel width (*deg*) | 11 | 11 | 11 | 11 | 11 | 11 |
| Kernel shape | Inverted diagonal | Diagonal | Diagonal | Diagonal | Diagonal | Diagonal |
| Synaptic probability[1] | 0.9 | 0.9 | 0.1 | 1 | 0.0001 | 0.05 |
| Maximum connection probability[2] | 0.438 | 0.371 | 0.9 | 0.9 | 0.9 | 0.9 |

[1]Fraction of connections in all possible pairs of neurons in pre- and postsynaptic populations.

[2]Probability that a neuron in the postsynaptic population receives at least one synapse from a neuron in the presynaptic population.

**Supplementary Table 4 - Spiking RNN best fitting model parameters.** Values of the main parameters of the best fitting model reported in Figure 5. A scaling factor was applied to $g_{max}$ to compensate for differences between model and experimental conditions.
